# Expression of *Wnt5a* defines the major progenitors of fetal and adult Leydig cells

**DOI:** 10.1101/2020.07.25.221069

**Authors:** Herta Ademi, Isabelle Stévant, Chris M Rands, Béatrice Conne, Serge Nef

## Abstract

Leydig cells (LCs) are the major androgen-producing cells in the testes. They arise from steroidogenic progenitors, whose origins, maintenance and differentiation dynamics remain largely unknown. Here, we identified *Wnt5a* as a specific marker of steroidogenic progenitors, whose expression begins at around E11.5-E12.5 in interstitial cells of the fetal mouse testis. *In vivo* lineage tracing indicates that *Wnt5a*-expressing progenitors are initially present in large numbers in the fetal testis and then progressively decrease as development progresses. We provide evidence that *Wnt5a*-expressing cells are *bona fide* progenitors of peritubular myoid cells as well as fetal and adult LCs, contributing to most of the LCs present in the fetal and adult testis. Additionally, we show in the adult testis that *Wnt5a* expression is restricted to a subset of LCs exhibiting a slow but noticeable clonal expansion, revealing hitherto unappreciated proliferation of fully differentiated LCs as a contribution to the adult LC pool.

## Introduction

Leydig cells are the major steroidogenic cells in the testes. They synthetize androgens that are essential for development of the Wolffian duct derivatives, masculinization of secondary sexual characteristics such as male external genitalia and spermatogenesis. In rodents, two distinct populations of LCs have been identified, one arising during the fetal period - henceforth referred to as fetal Leydig cells or FLCs - and the other after puberty referred to as adult Leydig cells, abbreviated ALCs (Chen et al., 2016; Chen et al., 2017; Teerds and Huhtaniemi, 2015).

During the process of male sex determination, Sertoli cells are the first somatic cell type to differentiate at around E11.5 in mice (Albrecht and Eicher, 2001; Gubbay et al., 1990; Hacker et al., 1995; Koopman et al., 1990; Lovell-Badge and Robertson, 1990), and are essential for directing the differentiation of FLCs (Barsoum and Yao, 2006; Habert et al., 2001). The dramatic increase in the number of FLCs between E12.5 and E15.5 occurs through the recruitment and differentiation of Leydig progenitor cells via paracrine factors, rather than by mitotic division of differentiated FLCs (Barsoum et al., 2009; Barsoum and Yao, 2010; Bitgood et al., 1996; Brennan et al., 2003; Byskov, 1986; Kerr and Knell, 1988; Migrenne et al., 2001). During the first postnatal days, FLCs involute gradually and are replaced by ALCs, although some may persist in the adult testis (Haider, 2004; Shima et al., 2015; Wen et al., 2016). ALCs are not derived from pre-existing FLCs but, like their fetal counterparts, arise from uncharacterized LC progenitors located in the testicular interstitium (Barsoum et al., 2013; Davidoff et al., 2004; Kilcoyne et al., 2014; Shima et al., 2012).

The origin of FLCs and ALCs has been heavily debated over recent decades, but has not been completely resolved. A synthesis of current models suggests the presence of several types of Leydig progenitors derived from various sources that have the potential to give rise to both FLCs and ALCs. Recent studies using *in vivo* cell lineage tracing revealed that FLCs and ALCs derive from two fetal sources: (i) gonadal progenitors that originate from the coelomic epithelium and express the markers NR5A1, ARX, PDGFRA and (ii) mesenchymal progenitor cells from the gonad/mesonephric border that migrate into the developing testis and express Nestin and NOTCH1 (DeFalco et al., 2011; Karl and Capel, 1998; Kumar and DeFalco, 2018; Rotgers et al., 2018). In parallel, some studies indicated that both FLCs and ALCs develop from the same precursor pool, but that some of these progenitors remain dormant until prepubertal development, at which point they develop into ALCs (Barsoum et al., 2013; Davidoff et al., 2004; Ge et al., 2006; Svingen and Koopman, 2013). Indeed, perturbation of FLC differentiation by constitutive activation of the hedgehog pathway resulted in an increased number of FLCs and a parallel reduction in the number of ALCs later in puberty suggesting a common pre-established pool of progenitors for both FLCs and ALCs in the fetal testis (Barsoum et al., 2013). Finally, recent findings using *in vivo* lineage tracing revealed that a significant proportion of ALCs originate from dedifferentiated FLCs in mice (Shima et al., 2018).

Considering coelomic epithelium-derived progenitors, we have recently shown by single-cell RNA sequencing (scRNA-seq) that there is initially a unique coelomic epithelium-derived *Nr5a1*^+^ progenitor cell population at E10.5, contributing to both supporting and steroidogenic lineages in XX and XY gonads (Stevant et al., 2019; Stevant et al., 2018). A subset of these multipotent cells maintain their progenitor state and, by E12.5, undergo gradual transcriptional changes, acquiring a steroidogenic fate by progressively expressing *Pdgfra* (platelet-derived growth factor receptor alpha) (Brennan et al., 2003; Eliveld et al., 2019; Ge et al., 2006), *Arx* (aristaless-related homeobox) (Miyabayashi et al., 2013), *Nr2f2* (nuclear receptor subfamily 2 group F member 2 or *Couptf2*) (Kilcoyne et al., 2014; Qin et al., 2008) and *Tcf21* (transcription factor 21, or *Pod1*) (Cui et al., 2004). Whether these fetal steroidogenic progenitors persist during development and into adult life is unclear. Similarly, questions remain unanswered concerning the fate of these progenitors over time and their ability to produce FLCs and ALCs, or both, during development and later in life. Here we used scRNA-seq and inducible cell lineage tracing experiments to address these questions.

## Results

### Identification of *Wnt5a* as a marker of gonadal steroidogenic progenitors

Using time-series scRNA-seq on XX and XY somatic *Nr5a1*^+^ cells spanning the sex determination process, we identified a single population of multipotent progenitors that give rise to both pre-supporting cells and potential steroidogenic precursor cells (**Figure 1A**) (Stevant et al., 2019; Stevant et al., 2018). Further expression analysis of this single-cell data revealed that the *Wnt5a* gene is specifically enriched in somatic progenitors when they commit to a steroidogenic fate at around E12.5 (**Figure 1B-E**). We confirmed by RNAscope *in situ* hybridization (ISH) in mouse fetal testes at E12.5 that *Wnt5a* mRNA expression is restricted to interstitial cells and exhibits a dorso-ventral gradient consistent with the coelomic epithelial origin of these progenitors (**Figure 1F**) (Chawengsaksophak et al., 2012). This suggests that *Wnt5a* could be used as a cell marker to label this population of steroidogenic progenitors and follow their fate and differentiation during the process of testicular development and into adulthood.

**Figure 1:**
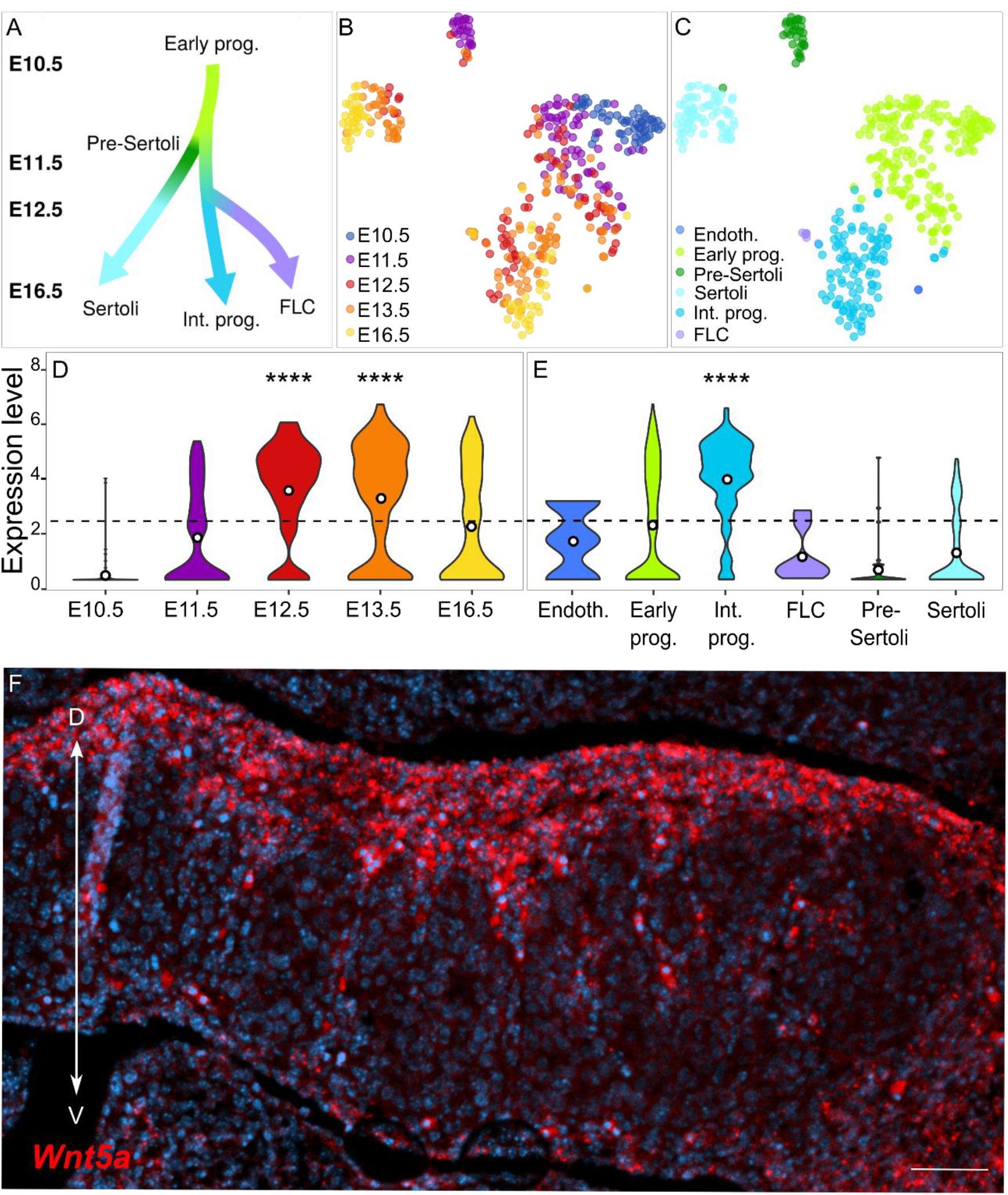
Identification of *Wnt5a* as a marker of the steroidogenic lineage and validation of its expression profile. (A) Schematic representation of the *Nr5a1* cell lineage at the origin of the supporting and steroidogenic lineages in XY gonads. (B) t-SNE representation of the 400 *Nr5a1*^+^ single cells from XY developing gonads colored by embryonic stages (B) and by cell clusters (C). *Wnt5a* expression represented as violin graph based on developmental stages (D) and *Nr5a1* cell cluster (E). The width of the “violin” indicates the proportion of the cells at that expression level. Expression scale presented as log (RPKM+1). The horizontal dashed lines represent the overall mean of expression, and the white dots in the violins are the mean of expression of the given cell populations. *Wnt5a* transcripts are highly expressed in the interstitial steroidogenic progenitors starting at E11.5 but not in the other branches of the *Nr5a1* lineage including early progenitors, pre-Sertoli cells, Sertoli cells and fetal Leydig cells. (F) RNA-Scope analysis revealed a dorso-ventral gradient of *Wnt5a* expression in the interstitial region of developing testis at E12.5. Abbreviations: Endoth., endothelial cells; Early prog., early progenitors; Int. prog., interstitial steroidogenic progenitors; FLC, fetal Leydig cells; Pre-Sertoli, pre-Sertoli cells; Sertoli, Sertoli cells. Difference of expression relative to the mean was assessed using the Wilcoxon test. ****P < 0.0001. D -- S, dorso-ventral axis. Scale bar (D) 50 μm.

### Use of the inducible Tet-On system for characterizing the fate of *Wnt5a*-expressing steroidogenic progenitor cells

To trace the contribution of the *Wnt5a*^+^ steroidogenic progenitors in the appearance of LCs in the developing testis and in adults, we employed an inducible Tet-On system. The Tet-On system allows non-invasive and efficient activation of a reporter gene by the administration of doxycycline (dox) in drinking water (Perl et al., 2002; Thorel et al., 2010). The use of doxycycline rather than tamoxifen in inducible systems to control Cre-mediated recombination has a major advantage for the study of reproductive organ development because tamoxifen exposure has been shown to cause adverse effects on the testis and the reproductive endocrine system (Patel et al., 2017). We first generated a *Wnt5a:Tet-On3G* knock-in line in which the tetracycline-inducible expression system transgene (Tet-On 3G) was inserted into the first coding exon of the *Wnt5a* gene (see Materials & Methods and **Figure 2A**), so that its expression accurately reflects the endogenous expression of *Wnt5a*. XX and XY heterozygous knock-in *Wnt5a:Tet-On3G* animals display no phenotype when compared to control animals (**Figure S1-S3** and **Supplementary data**).

**Figure 2:**
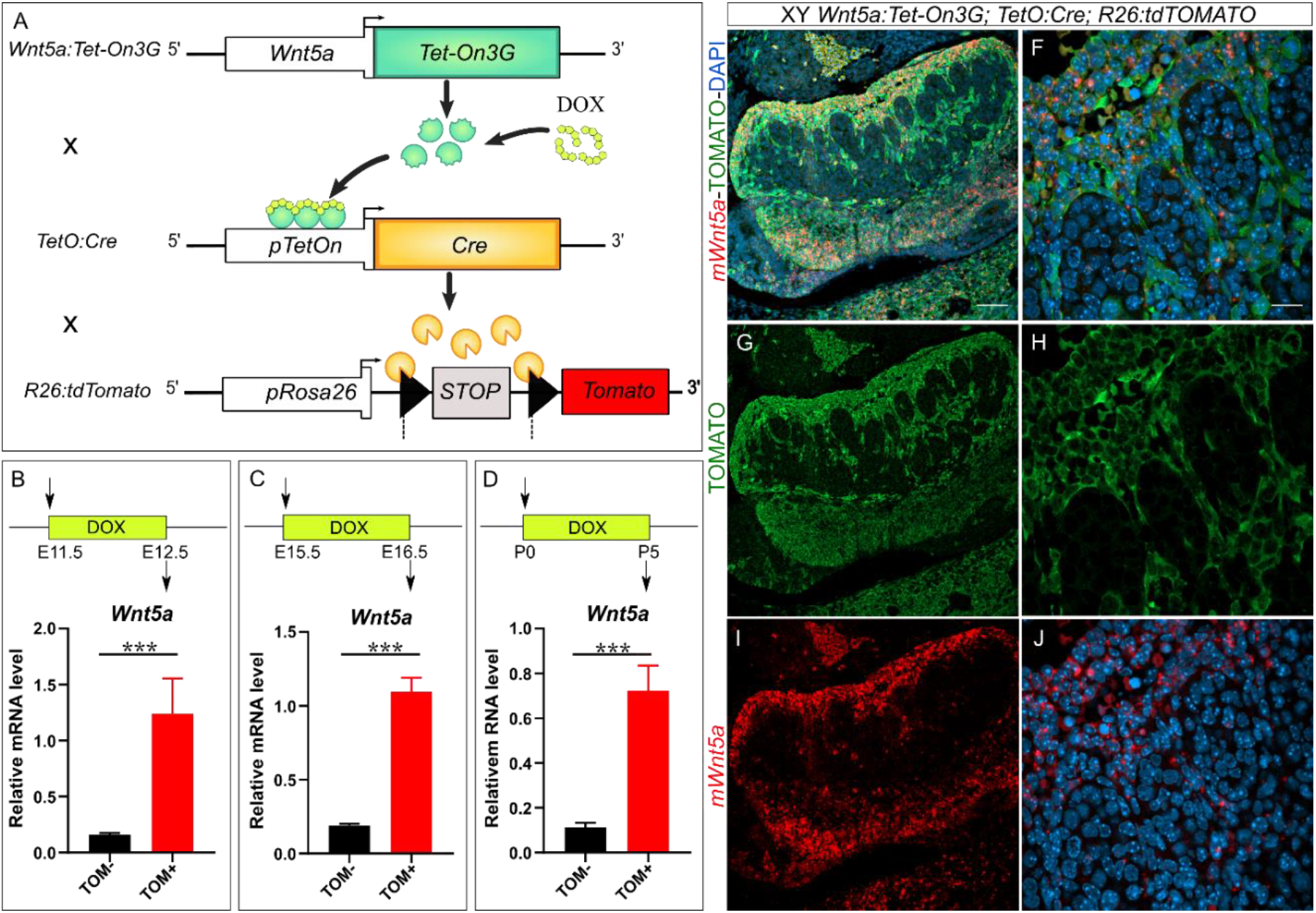
Lineage tracing strategy and assessment of the penetrance and specificity of the mouse model. (A) Schematic description of the lineage tracing strategy. Upon doxycycline (DOX) administration in the drinking water, *Wnt5a*-expressing cells of triple transgenic animals are permanently labelled by the Tomato marker. (B-D) Assessment of the efficiency and specificity of the labelling of *Wnt5a*-expressing cells. Quantitative RT-PCR performed with total RNAs from Tomato positive (+) or negative (−) cells isolated from triple transgenic testes. Dox labeling was induced either for 24 hours between E11.5 and E12.5 and the Tomato^+^ and Tomato^−^cells were isolated by FACs at E12.5 (n=3) (B), between E15.5 and E16.5 and the cells were isolated at E16.5 (n=3) (C), or between P0 and P5 and the cells were isolated at P5 (n=3)(D). Expression profiles were normalized to *Actin*, *Tubulin*, *Gapdh*. *** P <0.001. As expected, *Wnt5a* transcripts were highly enriched in all the Tomato^+^ cell populations. (E-J) RNA-Scope analysis for *Wnt5a* followed by immunostaining for the Tomato marker reveals co-expression in triple transgenic E12.5 testis dox-induced for 24 hours at E11.5. Nuclei were counterstained with 4′,6-diamidino-2-phenylindole (DAPI). Abbreviations: t, testis; m, mesonephros. Scale bar 50 μm.

We then crossed the *Wnt5a:Tet-On3G* knock-in line with *TetO:Cre* and *R26:tdTomato* reporter lines to generate triple transgenic *Wnt5a:Tet-On3G;TetO:Cre;R26:tdTomato* mice (**Figure 2A**). These mice enable dox-inducible, irreversible expression of tdTomato in *Wnt5a*-expressing cells. As expected, Tomato^+^ cells were not detected in both *Wnt5a:Tet-On3G;TetO:Cre;R26:tdTomato* without dox treatment and in those of control genotypes (i.e. *Wnt5a:Tet-On3G;R26:tdTomato* and *TetO:Cre;R26:tdTomato*) with dox induction. To confirm the high labeling efficiency of our *Wnt5a* cell lineage tracing system, we first verified by quantitative RT-PCR (qRT-PCR) that *Wnt5a* transcripts were specifically expressed in Tomato^+^ testicular cells. We chose this approach because no antibody worked effectively to detect WNT5A by immunofluorescence. For this purpose, *Wnt5a:Tet-On3G;TetO:Cre;R26:tdTomato* embryos were induced with dox at E11.5 for 24 hours. At E12.5 testes were dissected and Tomato^+^ and Tomato^−^ populations were sorted by fluorescence-activated cell sorting (FACS). We found that *Wnt5a* transcripts were highly enriched in the pool of purified Tomato^+^ cells, confirming that Tomato labelling is efficient and specific to cells expressing *Wnt5a* (**Figure 2B**). Similar results were obtained when Tomato labelling was induced by dox at E15.5 for 24 hours or at post-natal day (P) 0 for five days (**Figure 2C&D**). In addition, we performed *in situ* hybridization for *Wnt5a* transcripts and immunofluorescence (IF) for Tomato on 24-hour dox-induced E12.5 testis sections. In agreement with the qRT-PCR data (**Figure 2B**), the double labelling revealed strong *Wnt5a*-Tomato co-labelling and similar interstitial cell-restricted expression profiles with a dorso-ventral gradient (**Figure 2E-J**). All together, these results demonstrate that our *Wnt5a:Tet-On3G;TetO:Cre;R26:tdTomato* mouse model can be used for a specific and efficient dox-inducible, irreversible labelling of *Wnt5a*^+^ cells.

### *Wnt5a*^+^ cells dox-induced at E11.5-E12.5 are steroidogenic progenitors contributing to the majority of FLCs and ALCs

In order to study *in vivo* the presence of *Wnt5a*^+^ steroidogenic progenitors and their fate during testicular development from fetal age to adulthood, we performed a lineage tracing of the cells expressing *Wnt5a* in the fetal testis. Based on our knowledge of *Wnt5a* expression during early testis differentiation, E11.5 *Wnt5a:Tet-On3G;TetO:Cre;R26:tdTomato* embryos were dox-induced for a short period of 24 hours and testes were analyzed by immunofluorescence at E12.5, E13.5, P0, P10 and P60 (**Figure 3**). Whole-organ fluorescence analysis reveals strong Tomato signal in the interstitial compartment from E12.5 onwards (**Figure 3A-E**). At E12.5, we found that most Tomato^+^ cells co-expressed NR2F2 (also known as COUP-TFII) and/or ARX, indicating that *Wnt5a*-expressing cells display the characteristics of steroidogenic progenitors (**Figure 3F, F’, K**). At E13.5 and later stages, we observe a decreasing proportion of Tomato^+^ cells that co-express NR2F2 and ARX over time (**Figure 3 G-J, G’-J’, L-O**). It should be noted, however, that even in the adult testis (P60) a few Tomato and ARX co-expressing cells were still present, suggesting the persistence of a small pool of steroidogenic progenitors in adulthood (**Figure 3O**). To determine whether the *Wnt5a*^+^ steroidogenic progenitor cells also have a perivascular origin, we co-stained Tomato with Nestin, an intermediate filament protein expressed in many stem cell populations (Wiese et al., 2004), including a multipotent population of perivascular cells of mesonephric origin giving rise to LC, pericytes and smooth muscle cells (Jiang et al., 2014; Kumar and DeFalco, 2018). Our analysis of testes at E12.5 and E13.5 reveals some co-labelling, suggesting that some *Wnt5a*^+^ steroidogenic progenitors may have a perivascular origin (**Figure S4**).

**Figure 3:**
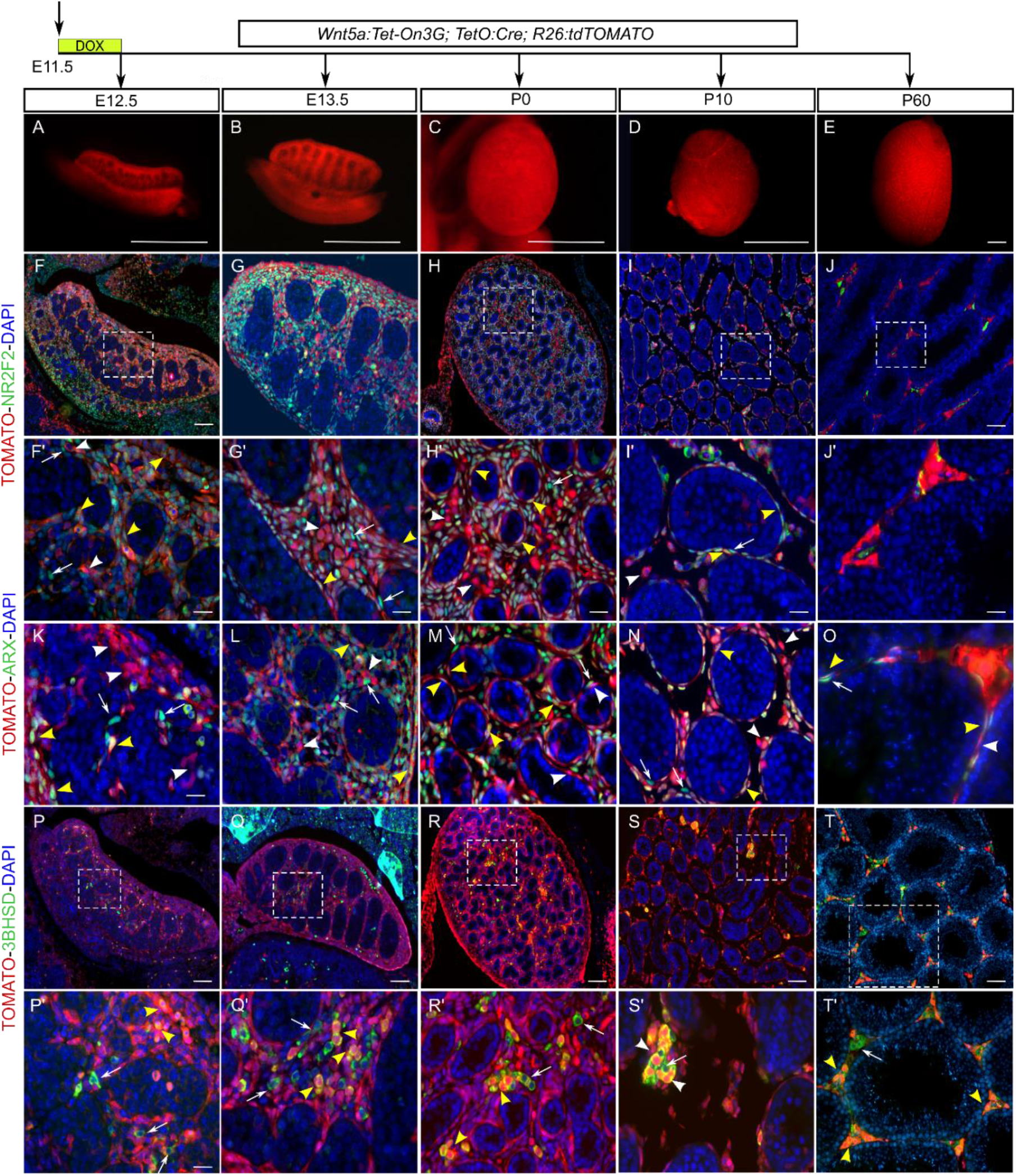
Lineage tracing of E11.5-E12.5 *Wnt5a*-expressing cells in developing and adult testes. (A-E) whole testis fluorescence of *Wnt5a:Tet-On3G; TetO:Cre; R26:tdTOMATO* animals dox-induced for 24h at E11.5. Note the high level of fluorescence in the interstitial compartment of the developing testis and in the mesonephros. Immunofluorescence image of triple transgenic testes at E12.5 (F,F’,K,P,P’), E13.5 (G,G’,L,Q,Q’), P0 (H,H’,M,R,R’), P10 (I,I’,N,S,S’) and P60 (J,J’,O,T,T’) exposed to dox between E11.5 and E12.5. Assessment of the identity of Tomato-labeled cells was obtained by double-immunofluorescence for Tomato and the progenitor markers NR2F2 (F-J and F’-J’), ARX (K-O) or the Leydig cell marker 3βHSD (P-T, P’-T’). During testis development, a large fraction of interstitial cells are Tomato-labelled, many of them co-expressing the progenitor markers NR2F2 and ARX. In the adult testis, only a small subset of these Tomato-labelled cells still express the ARX progenitor marker, and are localized closed to the seminiferous tubules (yellow arrowhead in O). From E13.5 onwards, there is a rapid increase in Tomato positive cells which also express the Leydig cell marker 3βHSD. At P10 and P60, the majority of the positive cells in 3βHSD are Tomato positive. Yellow arrowheads indicate co-expression, white arrowheads indicate Tomato expression, and white arrows indicate expression only in green channel. Nuclei were counterstained with 4′,6-diamidino-2-phenylindole (DAPI). Scale bars: 1mm (A-E), 10 μm (F’-O, P’-T’) 20 μm (F-G, P-Q),), 50 μm (I-J, S-T), 100 μm (H-R).

Considering LCs, few differentiated FLCs expressing the 3βHSD marker were detected at E12.5 (**Figure 3P, P’**). From E13.5 onward, we observed a rapid increase in the number of Tomato^+^ cells co-expressing the LC marker 3βHSD, in line with the idea that these progenitors give rise to most of the LCs in the fetal and adult testis (**Figure 3Q, Q’**). Quantitative analysis (**Table 1**) revealed that the proportion of double-labelled Tomato^+^/3βHSD^+^ FLCs increased from 28% initially at E12.5 to 56% at P0, indicating that the majority of FLCs come from *Wnt5a*-expressing progenitors specified between E11.5 and E12.5. Ultimately, at adult stages the *Wnt5a*-expressing cells labelled at E11.5-E12.5 represented nearly 71% of the total number of 3βHSD^+^ LCs.

**Table 1.** Quantitative data from Tomato-3βHSD double immunofluorescence analyses

| Doxycycline<br>induction | Analysis<br>Stage | Biological<br>replicates | Technical<br>replicates | Total<br>Tom<sup>+</sup><br>cells | Tom<sup>+</sup><br>only<br>cells | Total<br>3bHSD<sup>+</sup><br>cells | 3bHSD<sup>+</sup><br>only<br>cells | Tom<sup>+</sup>;3bHSD<sup>+</sup><br>cells | Tom<sup>+</sup>;3bHSD<sup>+</sup>/total<br>3bHSD<sup>+</sup> cells (%) |
| --- | --- | --- | --- | --- | --- | --- | --- | --- | --- |
| E11.5-P5 | P5 | 3 | 5 | 17762 | 15937 | 2199 | 374 | 1825 | 83.0 |
|  | P60 | 4 | 14 | 161633 | 100784 | 67749 | 6900 | 60849 | 89.8 |
| E11.5-E12.5 | E12.5 | 5 | 12 | 2771 | 2435 | 740 | 404 | 336 | 45.4 |
|  | E13.5 | 3 | 5 | 904 | 687 | 832 | 615 | 217 | 26.1 |
|  | P0 | 2 | 3 | 1545 | 1240 | 541 | 236 | 305 | 56.4 |
|  | P5 | 2 | 3 | 1013 | 904 | 297 | 188 | 109 | 36.7 |
|  | P10 | 1 | 2 | 11145 | 10450 | 794 | 99 | 695 | 87.5 |
|  | P20 | 1 | 3 | 8297 | 5413 | 4599 | 1715 | 2884 | 62.7 |
|  | P60 | 2 | 5 | 30277 | 19345 | 14842 | 3910 | 10932 | 73.7 |
| P0-P5 | P5 | 2 | 4 | 3414 | 3371 | 2252 | 2209 | 43 | 1.9 |
|  | P20 | 1 | 8 | 3786 | 3731 | 1386 | 1331 | 55 | 4.0 |
|  | P60 | 4 | 8 | 11013 | 6974 | 23745 | 19706 | 4039 | 17.0 |
| P60-65 | P65 | 3 | 15 | 641 | 43 | 114385 | 113787 | 598 | 0.5 |
|  | P85 | 4 | 14 | 1463 | 69 | 62698 | 61304 | 1394 | 2.2 |
|  | P180 | 3 | 12 | 5017 | 296 | 142031 | 137310 | 4721 | 3.3 |

Overall, this data suggests that the *Wnt5a*^+^ cells induced by dox between E11.5 and E12.5 initially possess an identity of steroidogenic progenitors. This early pool of *Wnt5a*^+^ steroidogenic progenitors decreases over time, although a small proportion persists in adulthood and gives rise to the majority of FLCs and ALCs present in the fetal and adult testes.

### *Wnt5a*^+^ cells dox-induced at P0-P5 are steroidogenic progenitors that give rise to 17% of ALCs present in the adult testis

To determine if *Wnt5a*-expressing cells are still present at early postnatal stages and what their fate is, *Wnt5a:Tet-On3G;TetO:Cre;R26:tdTomato* new born pups were dox-induced for a period of five days (P0-P5) and testes analyzed by immunofluorescence at P5, P20 and P60 (**Figure 4**). As expected, whole-organ fluorescence analysis revealed a robust signal restricted to the interstitial compartment of postnatal and adult testes (**Figure 4A-C**). At P5, we found that Tomato^+^ cells were localized in the interstitium and usually co-expressed the markers NR2F2 and ARX, indicating that *Wnt5a*-expressing cells are mostly steroidogenic progenitors (**Figure 4D-L**). At later stages, we observed a drastic decrease in cells co-expressing Tomato and ARX, although they were still present in low numbers, suggesting the persistence of a pool of steroidogenic progenitors in adulthood. For Leydig cells, double immunofluorescence analysis revealed that co-expression of the LC marker 3βHSD and Tomato at P5 was rare and these cells represented only 1.9% of 3βHSD^+^ cells (**Figure 4M-R** and **Table 1)**. These results are consistent with the observation that *Wnt5a* expression is restricted to steroidogenic progenitors and not LCs. In contrast, we observed that the proportion of LCs co-expressing Tomato increased to 4.0% at P20 and then to 17.0% in adult testis at P60 (**Figure 4N-O, Q-R** and **Table 1)**.

**Figure 4:**
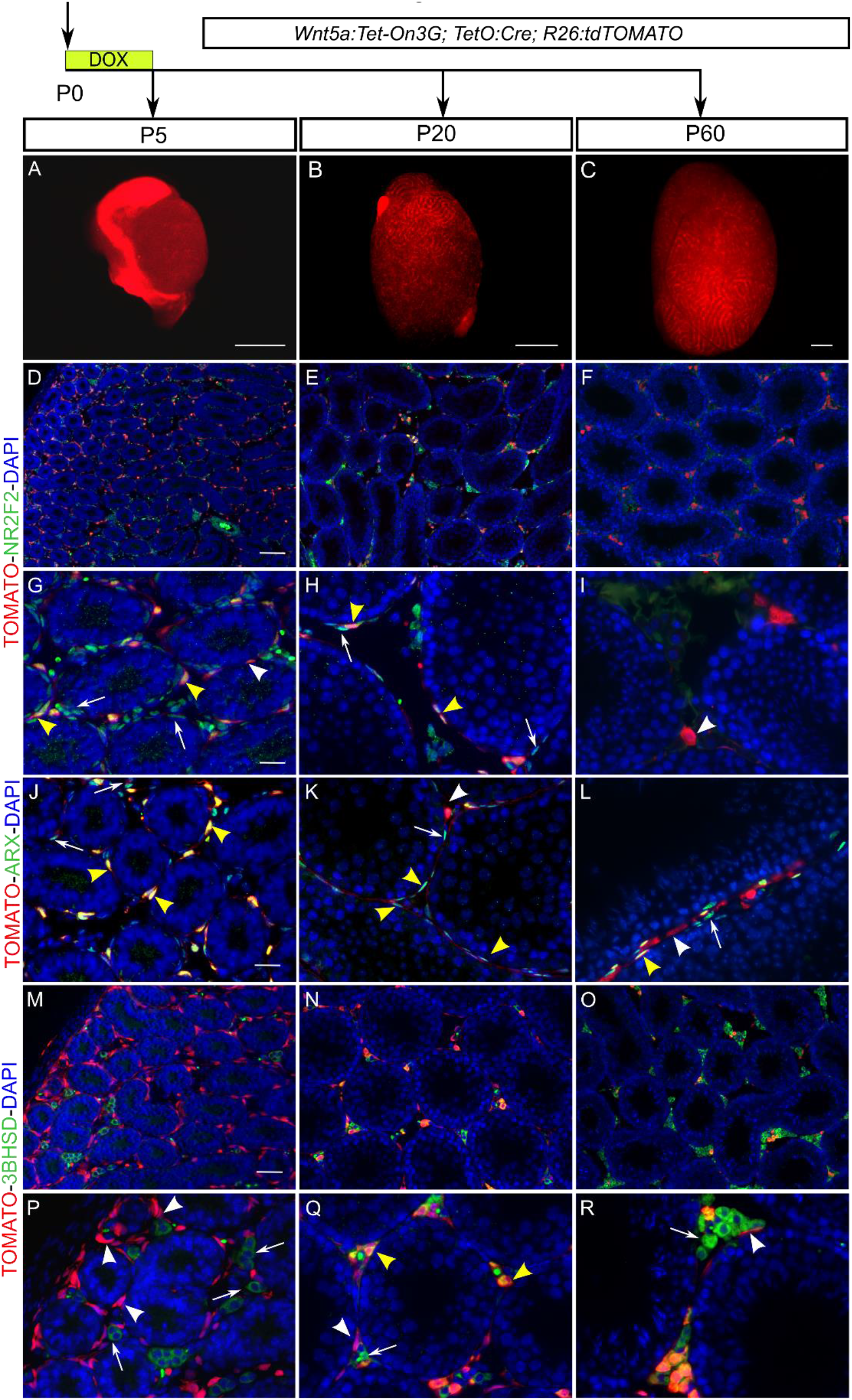
Lineage tracing of P0-P5 *Wnt5a*-expressing cells in developing and adult testes. (A-C) whole testis fluorescence of *Wnt5a:Tet-On3G; TetO:Cre; R26:tdTOMATO* animals dox-induced for 5 days at P0. Note the high level of fluorescence in the interstitial compartment of the developing testis. Immunofluorescence image of triple transgenic testes at P5 (D,G,J,M,P,S), P20 (E,H,K,N,Q,T) and P60 (F,I,L,O,R,U) exposed to dox between P0 and P5. Assessment of the identity of Tomato-labeled cells was obtained by double-immunofluorescence for Tomato and the progenitor markers NR2F2 (D-F and G-I), ARX (J-L) and the Leydig cell marker 3βHSD (M-O, P-R). At P5 and P20, most of Tomato-labelled cell co-express the progenitor markers NR2F2 and ARX. In the adult testis, only a small subset of these Tomato-labelled cells still express the ARX progenitor marker. At P5, the Leydig cells, as marked by 3βHSD were mutually exclusive for Tomato-lineage-traced *Wnt5a* expressing cells (M,P). By P20, a subset of Tomato-labeled cells expressed 3βHSD (N,O,Q,R). Yellow arrowheads indicate co-expression, white arrowheads indicate Tomato expression, and white arrows indicate expression only in green channel. Nuclei were counterstained with 4′,6-diamidino-2-phenylindole (DAPI). Scale bars: 1mm (A-C), 50 μm (D-F), 20 μm (M-O), 10 μm (G-L, P-R).

Overall, this data indicates that the *Wnt5a*^+^ cells induced by dox between P0 and P5 initially possess an identity of steroidogenic progenitors whose numbers decrease drastically over time, although a small proportion persist into adulthood. This pool of postnatal *Wnt5a*^+^ steroidogenic progenitors ultimately gives rise to a small proportion (17%) of ALCs present in the adult testes.

### A 12-day induction window between E11.5 and P5 reveals that *Wnt5a*^+^ cells give rise to ~90% of the LCs present in the adult testis

It remains unclear whether the pool of *Wnt5a*^+^ steroidogenic progenitors is generated within a short window of development at around E11.5-E12.5, or whether these cells appear over an extended period covering the second part of fetal development and the early postnatal period. To answer this question and evaluate the overall contribution of the *Wnt5a*^+^ progenitors to interstitial cells during testicular development, *Wnt5a:Tet-On3G;TetO:Cre;R26:tdTomato* embryos were dox-induced for an extended period of 12 days, from E11.5 to P5. Testes were then analyzed by immunofluorescence at P5 and P60, and the proportion of tomato-positive LCs was quantified (**Figure 5**). As expected, we found that Tomato^+^ cells were localized exclusively in the interstitium. More surprisingly, almost all interstitial cells were Tomato^+^. Concerning LCs, quantitative analysis of double immunofluorescence revealed that at P5 83% of the 3βHSD^+^ cells were also Tomato^+^. This number increased to 90% at P60, clearly indicating that *Wnt5a*-expressing cells are responsible for the vast majority of LCs, whether FLCs or ALCs. (**Figure 5** and **Table 1)**. Overall, these results suggest that *Wnt5a*^+^ cells induced by dox over a 12-day period between E11.5 and P5 ultimately give rise to most of the LCs present in both the fetal testis (83%) and the adult testis (90%).

**Figure 5:**
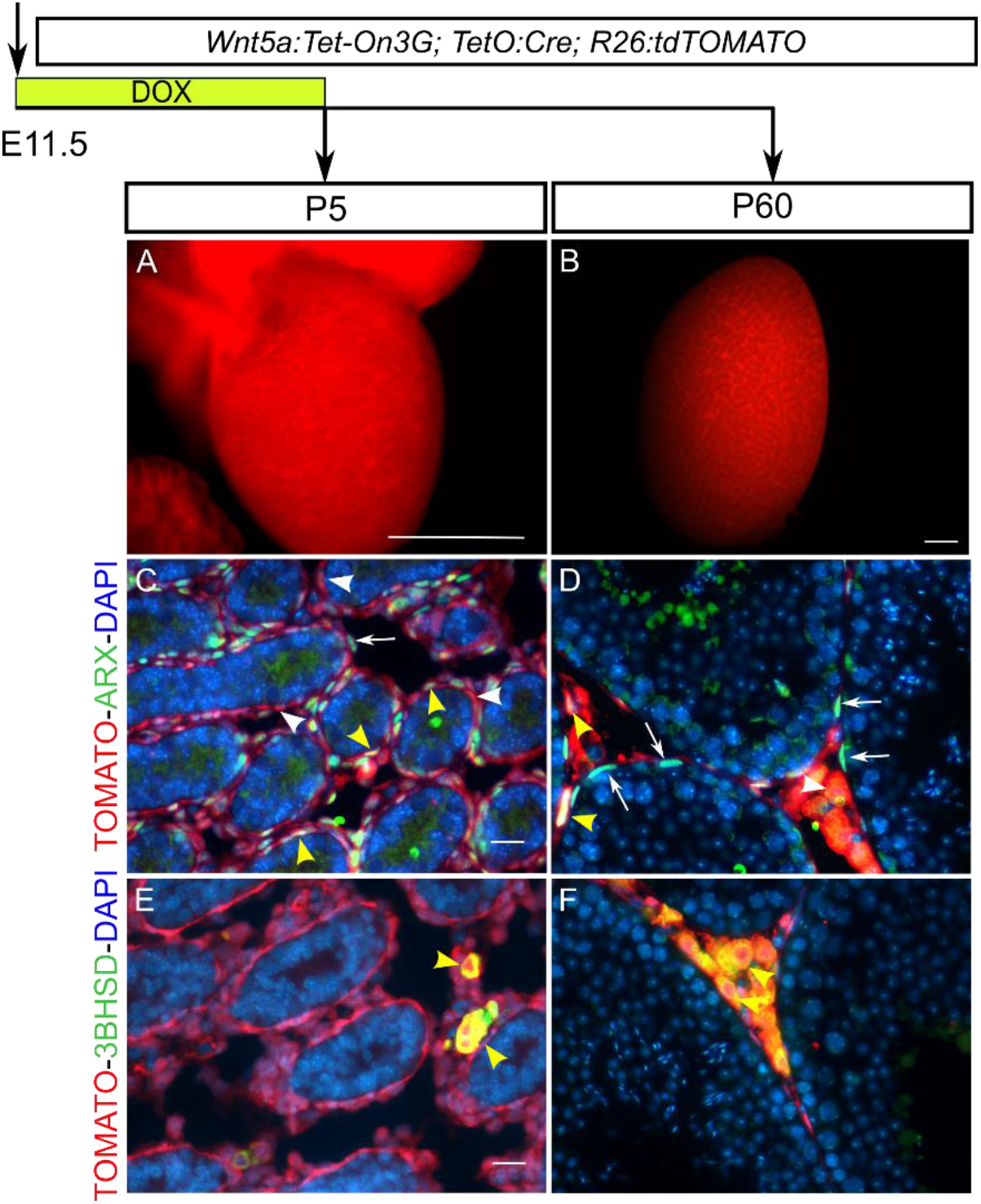
Lineage tracing of E11.5-P5 *Wnt5a*-expressing cells in post-natal and adult testes. (A-B) whole testis fluorescence of *Wnt5a:Tet-On3G; TetO:Cre; R26:tdTOMATO* animals dox-induced from E11.5 to P5. Note the high level of fluorescence in the interstitial compartment of the developing testis. Immunofluorescence image of triple transgenic testes at P5 (C,E,G) and P60 (D,F,H) exposed to dox between E11.5 and P5. Assessment of the identity of Tomato-labeled cells was obtained by double-immunofluorescence for Tomato and the progenitor marker ARX (C-D) and the Leydig cell marker 3βHSD (E, F). Yellow arrowheads indicate co-expression, white arrowheads indicate TOMATO expression, and white arrows indicate expression only in green channel. Nuclei were counterstained with 4′,6-diamidino-2-phenylindole (DAPI). Scale bars: 1mm (A-B), 10 μm (C-F).

### Fetal but not post-natal *Wnt5a*^+^ progenitors differentiate into peritubular myoid cells

To determine if *Wnt5a*-expressing cells give rise to other cell types in the developing testis, and in particular peritubular myoid cells (PMCs), we evaluated the co-expression of Tomato+ cells with ACTA2 (also called SMA), a marker of PMCs and vascular smooth muscle cells. To this end, we investigated the presence of double-positive cells for Tomato and ACTA2 in testes at P5 and P60, in animals whose *Wnt5a*-expressing cells were labelled by dox induction during three different time periods, namely E11.5-E12.5, E11.5-P5 or P0-P5. With regard to fetal induction (E11.5-E12.5 and E11.5-P5), we found that the vast majority of cells expressing the PMC marker ACTA2 co-expressed Tomato at both P5 and P60 (Figure 6A,B,D,E). In contrast, late induction at P0-P5 revealed a lack of co-labelling with ACTA2 (Figure 6C,F). These results indicate that the fate of *Wnt5a*^+^ progenitors is plastic and that only progenitors expressing *Wnt5a* during the fetal period are able to differentiate into PMCs.

**Figure 6:**
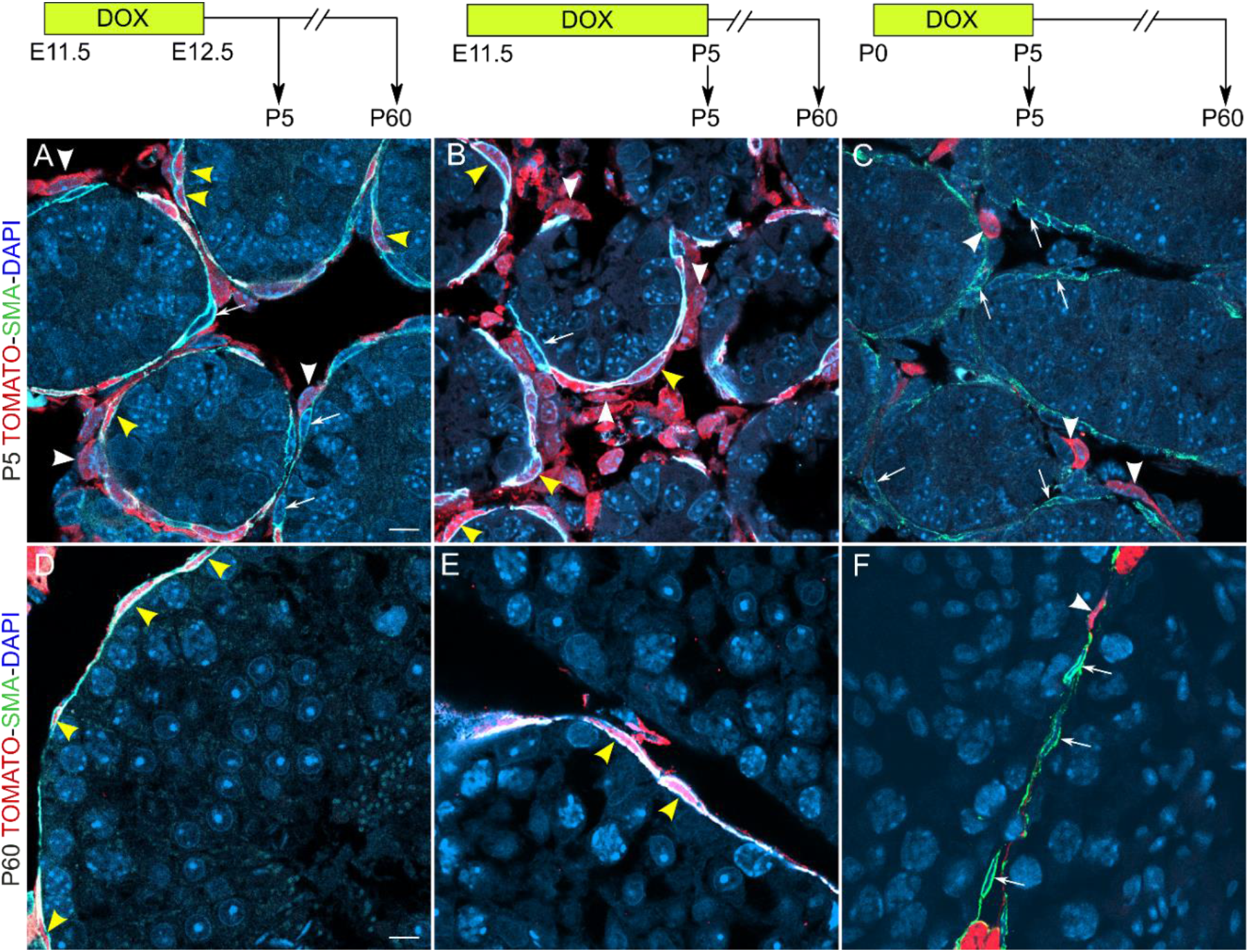
Co-immunofluorescence for tdTOMATO and ACTA2 in P5 and P60 testis induced by Dox at different developmental periods. Evaluation of Tomato-labelled cells and ACTA2 (SMA) expressing cells in P5 (A-C) and P60 (D-F) testes of *Wnt5a:Tet-On3G;TetO:Cre;R26:tdTOMATO* mice induced with Dox at E11.5-E12.5 (A,D) or E11-5-P5 (B,E) or P0-P5 (B,C,F). ACTA2-Tomato co-labelling is observed only with fetal inductions and is absent in perinatal induction (P0-P5). Yellow arrowheads indicate co-expression, white arrowheads indicate TOMATO expression. Scale bars: 10 μm (A,D).

### *Wnt5a*^+^ adult Leydig cells are proliferating at a low rate

In order to evaluate whether *Wnt5a* is still expressed in the adult testis and if so in which cell type, we analyzed scRNA-seq data from adult mouse testis (see (Ernst et al., 2019) and Material & Methods). Although the majority of testicular cells in this transcriptomic atlas are composed of germ cells at different stages of spermatogenesis, we identified a group of 828 ALCs based on the expression of the markers *Insl3* and *Hsd3b1* (which encodes 3βHSD) (**Figure 7A,B**& **D,E**). *Wnt5a* is expressed at detectable levels only in ALCs and, more specifically, in a subset of ALCs (**Figure 7C**& **F**). Further analysis suggests no substantial transcriptomic differences between these two (*Wnt5a*^+^ and *Wnt5a*^−^) populations, indicating that *Wnt5a*^+^ ALCs are likely to be representative of ALCs as a whole.

**Figure 7:**
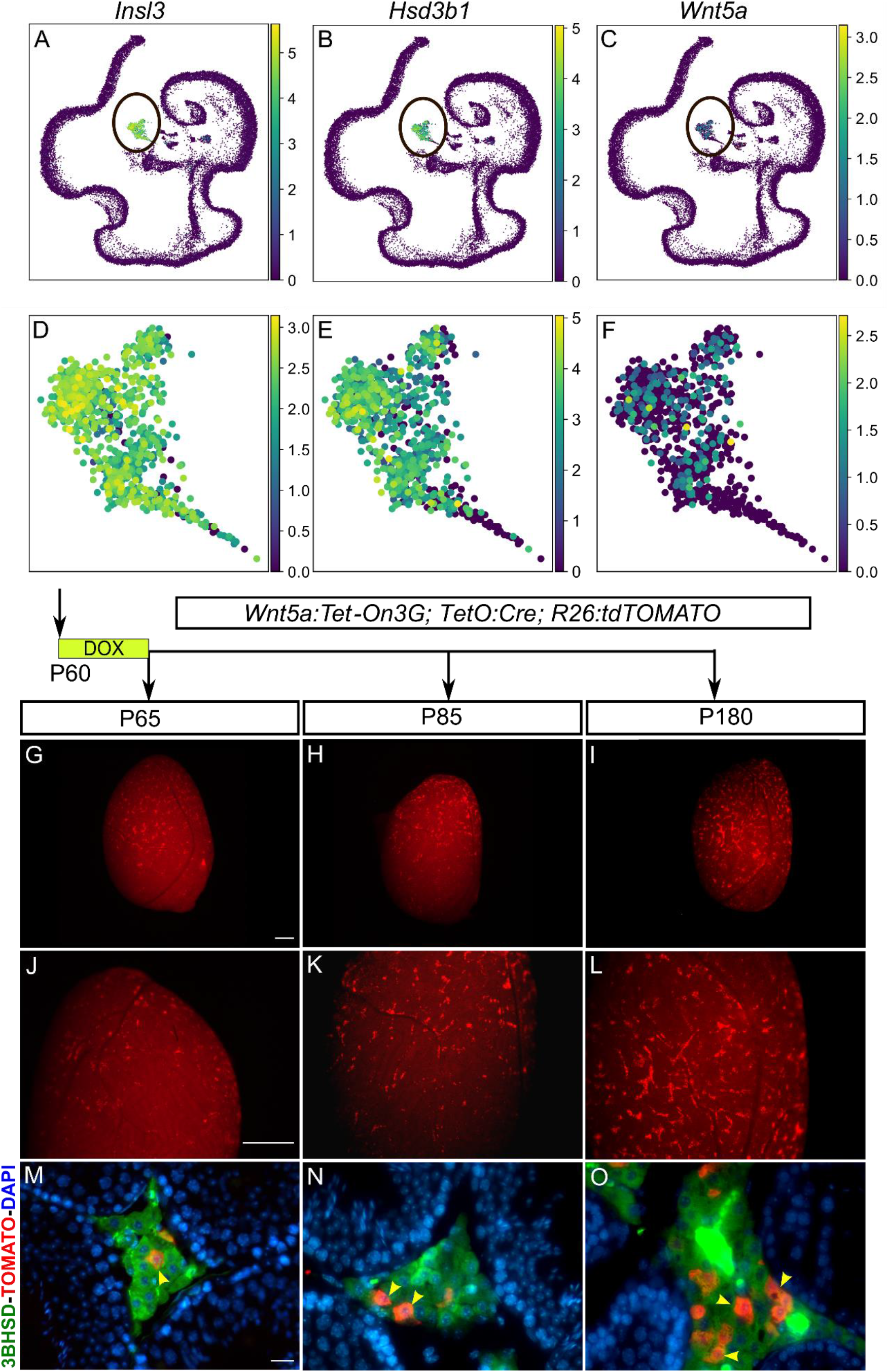
*Wnt5a* is expressed in a subpopulation of adult Leydig cells at P60 that increases in number over time. 10X scRNA-seq data of adult (age 8-10 weeks) mouse testes from Ernst et al. 2019 was processed (see methods) and LC cluster was identified with the expression of *Insl3* and *Hsd3b1* (A-B). (C) *Wnt5a* expression profile in the whole P60 testes is restricted to the LC cluster. (D-F) Zoomed view of *Insl3, hsd3β1, Wnt5a* expression within LC cluster, where *Wnt5a* expressing cells are homogenously distributed in the group. (G-O) Lineage tracing of *Wnt5a*-derived LCs in *Wnt5a:Tet-On3G; TetO:Cre; R26:tdTOMATO* adults induced for 5 days from P60 to P65. (G-L) Whole testis fluorescence of triple transgenic animals dox-induced as observed at P65, P85 and P180. Note the rare initial fluorescent labelled cells scattered in the interstitial compartment of the testis at P60, whose number increase towards P180. Evaluation of the identity of Tomato-labeled cells at P65 (M), P85 (N) and P180 (O) by co-immunofluorescence for tdTOMATO and the Leydig cell marker 3βHSD. Not only do my Tomato and 3βHSD tags co-locate but the density of double positive cells increases with time suggesting clonal multiplication of ALCs. Yellow arrowheads indicate co-expression. Nuclei were counterstained with 4′,6-diamidino-2-phenylindole (DAPI). Scale bars: 1mm (G-L), 10 μm (M-O).

To study the fate of *Wnt5a*^+^ ALCs in the adult testis, *Wnt5a:Tet-On3G;TetO:Cre;R26:tdTomato* adult males were induced with dox for 5 consecutive days at P60. At P65, whole-organ fluorescence analysis revealed rare and scattered signals in the testicular interstitium (**Figure 7G**& **J**). The signal profile is similar after 20 days and 4 months (at P85 and P180, respectively), although the positive cells appear more frequently and are often clustered at the later stage (**Figure 7H,I**& **K,L**). A double labelling by immunofluorescence for Tomato and 3βHSD reveals that at P65, the rare Tomato^+^ cells detected were also 3βHSD^+^ and located in the interstitial compartment (**Figure 7M**). Consistent with our scRNA-seq expression data, these cells display all the characteristics of ALCs. Taking advantage of the rare and scattered Tomato^+^ labelling of ALCs present in the adult testis at P65, we decided to use it to trace the fate of these cells. While at P65, Tomato^+^ ALCs were rare and isolated, at P180 we observed Tomato^+^ ALCs located in clusters, usually near blood vessels, suggesting clonal expansion (**Figure 7N, O**). Quantitative analysis of immunofluorescence data indicated that the fraction of double Tomato- and 3βHSD-labelled ALCs in physiological conditions increased from 0.5% at P65 to 2.4% at P85 and finally 3.3% at P180, reflecting an increase in the proportion of Tomato^+^ ALCs of more than 6-fold (see **Table 1**). Clonal proliferation will result in the presence of clusters of Tomato^+^ cells whose constituent cells will be very close to each other. If this is indeed the case, we should observe a reduction in the mean distance between Tomato^+^ cells between P65 and P180 as a consequence of clonal proliferation. As predicted, spatial analysis revealed that the mean distance between Tomato^+^ ALCs decreased significantly from 242μm at P65 to 164μm at P85 and 79μm at P180 (F(2.39)=4.16, p=0.023) (**Figure 7O**). A post-hoc analysis based on Tukey’s multiple comparison test showed a significant difference between P65 and P180 (p=0.0172). Overall, these results suggest that *Wnt5a*^+^ ALCs physiologically proliferate at a slow pace in the adult testis.

## Discussion

Understanding the steroidogenic lineage(s) through which testosterone-producing cells are formed is a fundamental question in developmental and reproductive biology. Several technical challenges are associated with the study of steroidogenic progenitors in developing gonads. First, early specific markers are scarce, usually not specific and rarely linked with function. In addition, these steroidogenic progenitors operate in a dynamic environment influenced by intrinsic factors and environmental cues that alter their transcriptome and cell fate. In the present study, we combined single cell transcriptomics with inducible genetic lineage tracing to study the steroidogenic cell lineage in male mice, dissecting the developmental dynamics of steroidogenic progenitors under physiological conditions in fetal, perinatal and adult environments. Overall, our observations support the idea that *Wnt5a* expressing cells are *bona fide* LC progenitors and give rise to the majority of FLCs (80%) and ALCs (90%). We found that the fate of these progenitors varies according to the stage at which these cells express *Wnt5a*. More precisely, in the course of embryonic and postnatal testis development, the *Wnt5a* lineage potential become increasingly restricted. While fetal *Wnt5a*-expressing progenitors will give rise to FLCs, ALCs and PMCs, those expressing *Wnt5a* post-natally mostly differentiate into ALCs. Finally, in the adult testis, *Wnt5a* expression is restricted to a small subset of LCs. *In vivo* lineage tracing revealed that these cells proliferate at a low rate and potentially serve to replenish and maintain the LC population in the adult testis.

### The use of the non-canonical *Wnt5a* as a marker of steroidogenic progenitor lineage

WNT5A, a member of the wingless-related MMTV integration site (WNT) family, functions as a secreted morphogen and is associated with a wide range of developmental processes including differentiation and proliferation (He et al., 2008; Huang et al., 2009; Kim et al., 2005; Serra et al., 2011; Warr et al., 2009; Yamaguchi et al., 1999). In the developing testis, *Wnt5a* is expressed in interstitial cells of the differentiating testis, starting at around E11.5-E12.5 in a dorso-ventral gradient (see (Chawengsaksophak et al., 2012) and **Figure 1)**. Consistent with its expression profile, our findings confirm that *Wnt5a*^+^ cells are true steroidogenic progenitors giving rise to most FLCs and ALCs. Other genes expressed in steroidogenic progenitors might also have been potential candidates to use for genetically tracing the lineage of these cells, including *Nr5a1*, *Nr2f2*/*CouptfII*, *Pdgfra* and *Tcf21*/*Pod1* (Brennan et al., 2003; Shen et al., 2020). However, the expression of these genes is not restricted to the steroidogenic progenitors and they usually display a more widespread expression, either at earlier stages of development and/or in other cell types. Consequently, *Nr5a1*^+^, *Tcf21*^+^, *Nr2f2*^+^, *Nestin*^+^ or *Pdgfra*^+^ cells contribute to numerous somatic cell types in the testis and are not restricted to the steroidogenic lineage. *Tcf21*, encodes a basic helix-loop-helix transcription factor that is expressed in somatic cells of the bipotential gonadal ridge and is known to be important for testis development (Barsoum et al., 2009; Cui et al., 2004; Lu et al., 2000; Lu et al., 2002; Quaggin et al., 1999). Comparison of *Tcf21* and *Wnt5a* expression in the *Nr5a1*^+^ somatic cell lineage using scRNA-seq data revealed that *Tcf21* is expressed as early as E10.5 in *Nr5a1*^+^ progenitors, a stage when these cells are still multipotent and can differentiate into both the supporting and steroidogenic cell lineage (**Supplementary Figure S5**). In contrast, *Wnt5a* expression is initiated at a later stage (E12.5). This difference in expression explains why early *Tcf21*^+^ progenitors contribute to all known somatic population of the testis such as Sertoli cells, PMCs and LCs (Shen et al., 2020), while the fate of *Wnt5a*^+^ progenitors is mainly restricted to FLCs and ALCs, as reported here.

### *Wnt5a*^+^ cells are steroidogenic progenitors giving rise to the large majority of FLCs and ALCs

At least two different sources of steroidogenic progenitors give rise to FLCs and ALCs: the *Nr5a1+* CE-derived and the *Nestin*^+^ perivascular-derived progenitors. Our analysis by scRNA-seq focused only on the *Nr5a1* lineage. However, it may be possible that *Wnt5a* is also express in *Nestin*^+^ progenitors and may represent a more global marker for all classes of steroidogenic progenitors. Consistent with this hypothesis, we found that about 80% of the FLCs and 90% of the ALCs are derived from the *Wnt5a*^+^ progenitors when dox-induction is extended for a long period (E11.5 to P5). *In vivo* lineage tracing of *Nestin*^+^ progenitors revealed that ~33% of FLCs and ~50% ALCs were derived from these *Nestin*^+^ progenitors (Kumar and DeFalco, 2018). Taken together, our findings suggest that *Wnt5a* is a defining steroidogenic progenitor marker expressed in both *Nr5a1+* CE-derived and *Nestin*^+^ perivascular-derived progenitors. This hypothesis is supported by our RNAscope *in situ* hybridization data from E12.5 embryos, which clearly indicated the presence of *Wnt5a* transcripts in both the mesonephros and the interstitial compartment of the testis (Figure 2I).

### The fate of *Wnt5a*^+^ progenitors is plastic and varies according to the stage at which these cells express *Wnt5a*

Our results revealed that the fate of *Wnt5a*^+^ progenitors varies depending on the developmental stage at which these cells express *Wnt5a*. The *Wnt5a* progenitors may either maintain their progenitor status or differentiate into LCs or PMCs. This is particularly noticeable when we compare the fate of Tomato^+^ following dox induction performed in different developmental windows. While progenitor cells expressing *Wnt5a* during the fetal period give rise to LCs and PMCs, cells expressing *Wnt5a* in the early postnatal stage (P0-P5) differentiate almost exclusively into LCs. Thus, only progenitors expressing *Wnt5a* between E11.5 and P0 are able to differentiate into PTM cells (see **Figure 6**). This suggests that *Wnt5a* expressing cells are initially multipotent with several potential fates such as remaining progenitors or differentiating into FLCs, ALCs or PMCs, then later the fate is restricted to give almost exclusively ALCs. The findings that PMCs are largely derived from *Wnt5a*^+^ progenitors is not entirely surprising, as other recent lineage tracing studies investigating the fate of steroidogenic progenitors using markers such as *Nestin* or *Tcf21* obtained similar results (Kumar and DeFalco, 2018; Shen et al., 2020). In addition, another evidence indicative of a common progenitor is the role of DHH, a paracrine factor secreted by Sertoli cells, which is responsible for the differentiation of both PMCs and LCs (Pierucci-Alves et al., 2001; Yao et al., 2002). *Dhh*-null mice develop abnormal peritubular myoid and Leydig cells. However, it is currently unclear how this single multipotent steroidogenic progenitor is able to commit to a PMC or FLC fate and what testicular factors favor one fate over the other.

### *Wnt5a*^+^ Leydig cell undergo rare events of cellular divisions in adult testis

In adult testes, our scRNAseq and genetic lineage tracing results clearly indicate that *Wnt5a* is expressed in a small fraction of the ALCs. We also found that these *Wnt5a*^+^ ALCs undergo rare events of cell division in adult testis. This is particularly surprising because the current dogma clearly states that LCs do not proliferate, but instead originate from steroidogenic progenitor cells that proliferate and then differentiate into FLCs or ALCs (Hardy et al., 1989; Miyabayashi et al., 2013; Vergouwen et al., 1991). Our analyses based on scRNA-seq data did not reveal obvious transcriptional differences between *Wnt5a*^+^ and *Wnt5a*^−^ALCs populations, suggesting that ALCs are a homogenous population. To our knowledge, the only data suggesting potential renewal of LCs in the adult testis come from a study by Teerds et al (Teerds et al., 1989), which examined the renewal of Leydig and other interstitial cells in the adult rat testis by injecting 3H-thymidine twice daily for 1, 3 and 8 days. The percentage of labelled ALCs, which was initially low (0.5%), gradually increased during treatment to 1.4%. Our data are consistent with data from Teerds et al. and suggest that *Wnt5a*^+^ ALCs contribute to steroidogenic cell numbers in adult testes. Alternatively, it remains a theoretical possibility that *Wnt5a*^+^ ALCs de-differentiate into progenitor cells under physiological conditions in order to be able to reenter into the cell cycle, expand the progenitor pool and subsequently re-differentiate into ALCs. In both cases, this slow renewal process is remarkable, as it provides evidence of LC regeneration under physiological conditions. The fact that differentiated testosterone-producing cells can proliferate could have far-reaching implications in understanding and possibly treating patients with primary hypogonadism or late hypogonadism, where androgen biosynthesis is defective due to impaired LC function.

## STAR Methods

**Key Resources Table** (see attached document)

### Contact for Reagent and Resource Sharing

Further information and requests for resources and reagents should be directed to and will be fulfilled by the Lead Contact, Serge Nef.

### Method details

#### Animals

*(tetO)7-CMV-Cre (TetO:Cre)*, and *Gt(ROSA)26Sor*^*tm9(CAG-tdTomato)Hze*^ (*R26:rdTomato*) transgenic animals were described previously (Madisen et al., 2010; Perl et al., 2002). The *Wnt5a*^*tm1(Tet-On 3G)Nef*^ strain was obtained by inserting the 1.2kb *Tet-On3G* sequence 8 nucleotides after the ATG and 0.8 kb downstream of the *Wnt5a* initiation of transcription using the CrispR/Cas9 system (see **Sup. Material & Methods** for more details). Routine genotyping of the transgenic lines was performed by classic PCR using sets of primers specific for the *TetO-Cre* and *R26:tdTomato* transgenes as described previously, or a set of primers for *Wnt5a*^*tm1(Tet-On 3G)Nef*^ genotyping (see **Sup. Material & Methods**). To analyze the fate of *Wnt5a*-expressing cells, we performed lineage tracing using *Wnt5a*^*tm1(Tet-On3G)Nef*^;*(tetO)7-CMV-Cre*^*TG/+*^;*R26*^*tdTOMATO/+*^ mice later abbreviated *Wnt5a:Tet-ON3G;TetO-Cre;R26:tdTomato* for simplicity. *Wnt5a*^*Tet-On 3G/+*^;*TetO-Cre*^*TG/+*^;*R26*^*tdTOMATO/+*^ were considered as positive whereas *Wnt5a*^*Tet-On3G/+*^;*TetO-Cre+/+*;*R26*^*tdTOMATO*/+^, Wnt5a+/+;*TetO-CreTG/+*;*R26*^*tdTOMATO*/+^ and *Wnt5a^Tet-On 3G/+^*;*TetO-Cre*^*TG/+*^;*R26*^*tdTOMATO+/+*^ were considered as negative controls. DOX was administered in drinking water (pH 6.5) at a concentration of 2 mg/mL. Animals were then sacrificed at different time points to monitor the fate of *Wnt5a* expressing cells during testicular development and in adulthood. The animals were treated with care and respect. This study complies with all ethical rules applicable to animal research and all experiments were approved and conducted in accordance with the guidelines of the Service de la consommation et des affaires vétérinaires du Canton de Genève (licence numbers GE/67/19 and GE/105/18).

#### Single cell RNA sequencing and analysis

Single-cell RNA sequencing for *Nr5a1*^+^ cells in fetal gonads was generated as described in (Stevant et al., 2019; Stevant et al., 2018). Briefly, somatic cells of developing XY mouse gonads were purified by sorting gonadal cells from Tg(*Nr5a1:GFP*) embryos at 5 stages of development (E10.5, E11.5, E12.5, E13.5, and E16.5) using FACS BD ARIA II. Single-cell isolation, reverse transcription and cDNA amplification was performed using the Fluidigm C1 Autoprep system on the 96 well IFC chips, and single-cell sequencing libraries were prepared with Illumina Nextera XT following the Fluidigm protocol. Libraries were multiplexed and sequenced on an Illumina HiSeq2000 platform with 100bp paired-end reads, at an average depth of 10 million reads per single-cell. Obtained reads were mapped on the mouse reference genome (GRCm38.p3) and gene expression normalized and quantified in RPKMs (Reads Per Kilobase of exon per Million reads mapped). Cells were clustered with R using PCA and hierarchical clustering, and cell types were identified according to the expression level of marker genes and gene ontology enrichment tests.

10X scRNA-seq data of adult (age 8-10 weeks) mouse testes from Ernst et al. 2019 was pre-processed with CellRanger Count to map the sequencing reads to the GRCm38 mouse reference genome with GENCODE M15 transcriptome annotation (Ernst et al., 2019). This resulted in a filtered cells vs. genes matrix. Using Scanpy (Wolf et al., 2018), cells with fewer than 50 genes expressed and genes expressed in fewer than 3 cells were filtered out. Expression counts were normalized and log-transformed. Highly variable genes were identified. The top 50 PCA components were embedded in the neighbourhood graph with batch correction via BBKNN (Polanski et al., 2020) and the data was visualized with 2D UMAPs (McInnes et al., 2018). Cell clusters were defined with the Leiden algorithm (Traag et al., 2019) and the population of LCs was identified based on the expression of the classical marker genes *Insl3* and *Hsd3b1*. Computations were performed on the Baobab HPC cluster at the University of Geneva.

#### Immunostainings and RNAscope® in situ hybridization

Mouse embryos at E12.5 were collected and fixed overnight in 4% paraformaldehyde, serially dehydrated, embedded in paraffin and sectioned at 5-μm. Co-localization of *Wnt5a* transcripts and Tomato protein was performed with the RNAscope® 2.5 LS Assay-RED kit (ACDBio) according to manufacturer’s instructions for soft tissues. The RNAscope probe used in this study is *Mm-Wnt5a* (ACDBio). Images were acquired with ZEISS LSM800 confocal microscope equipped with a plan-apochromat 10x 0.45NA objective (0.624μm/pixel) or with a 40x 1.4NA oil immersion objective (0.156μm/pixel).

#### Flow cytometry, RNA extraction and quantitative RT-PCR

XX and XY gonads of *Wnt5a:Tet-On3G;TetO:Cre;R26:tdTOMATO* animals were collected at E12.5, E16.5 and P5. Testis or ovaries were dissociated in trypsin-EDTA 0.05% at 37°C for up to 10 minutes under rapid agitation. Cell suspensions were re-suspended in 0.5ml of DPBS and filtered through a 70-μm cell strainer (BD Falcon; BD Biosciences Discovery Labware, Bedford, Massachusetts) for fluorescent activated sorting (FACS) using a Cell sorter S3 (Bio-rad, Berkeley, California). The gating strategy consisted in debris exclusion (SSC vs FCS), dead cell exclusion (FCS vs Draq7 fluorescence intensity) and doublet exclusion (FSC height vs FSC area). Finally, Tomato-negative and - positive cell fractions were gated using the intensity on the FL1 channel. In E12.5 males, we detected on average 60% of Tomato^+^ cells, whereas only 40% and 5% were detected at E16.5 and P5, respectively.

For RNA extraction, pools of 40.000 FACS-sorted Tomato^+^ or Tomato^−^cells were extracted using an RNeasy MicroKit (Qiagen, Valencia, California), according to the manufacturer’s protocol. RNA integrity and quantity were assessed using RNA 6000 picochips with a 2100 bioanalyzer (Agilent Technologies, Santa Clara, California) before and after genomic DNA removal with DNaseI digestion. 500pg to 1ng of total RNAs was reverse-transcribed with random hexamers using the M-MLV reverse transcriptase (Promega, Madison, USA). A selection of relevant genes (*Wnt5a, Nr2f2, Arx, Amh, Cyp11a1, Actb, Tuba1a, Gadph*) were pre-amplified with the Prelude PreAmp Master Mix (Takara, Shiga, Japan). Real-time RT-PCR was performed on each sample in triplicate using KAPA SYBRFast (Kapa Biosystems, Wilmington, MA, USA) and a Corbett Rotor-Gene 6000 (Qiagen, Valencia, California). Data was analyzed using a comparative critical threshold (CT) method with the amount of target gene normalized to the average of 3 endogenous control genes (*Actb* and *Tuba1a* and *Gadph*). The primers used for qRT-PCR are listed in **Table S4**.

#### Histological and immunological analyses

Embryos were fixed overnight at 4°C in 4% paraformaldehyde, dehydrated in an ethanol series and embedded in paraffin. Five μm-sections were stained with hematoxylin and eosin (H&E) or processed for immunofluorescence (IF). For IF analysis, antigen retrieval was performed either in TEG buffer (10 mM, pH 9) or sodium citrate buffer (10 mM, pH 6) for 15 minutes in a pressure cooker. Sections were incubated in blocking reagent (3% bovine serum albumin, 0.1% tween in PBS) for 2 hours at room temperature, and primary antibodies incubated overnight at 4°C. Slides were counterstained with DAPI and mounted with PBS:glycerol (1:1). Fluorescent images were acquired using Axio Scope.A1 microscope fitted with an AxiocamMRc camera (Carl Zeiss, Oberkochen, Germany) and processed using the Zen software (Carl Zeiss).

#### Cell counting

To determine the percentage of positive cells for Tomato, ARX, 3βHSD as well as the double labeled Tomato/ARX or Tomato/3βHSD cells, we scanned the IF sections. In short, fluorescent images of double IF (Tomato/ARX and Tomato/3βHSD) were acquired with a slide scanner Axio Scan.Z1 (ZEISS, Oberkochen, Germany) using a Plan-Apochromat 20x/0.8 objective (0.325μm/pixel). Whole testis images were analyzed using QuPath v0.2.0 (Bankhead et al., 2017) to extract fluorescent intensities within all the detected cells. The cellular information was then processed with MATLAB R2019b (The Mathworks, Natick, Massachusetts, USA), to classify cells into red and green positive ones if their respective fluorescent intensities and homogeneities were above user defined thresholds. Then the variation of red and green positive cells through time was expressed as a percentage of positive cells within the population. For all experiments and stages, three to five different sections, from at least three independent gonads from different animals were used. Detail of total cell counts are included in **Table 1**. Subsequently, for the 3βHSD and Tomato experiments, the proximity was assessed based on the pairwise Euclidean distance between pairs of all Tomato positive cells. The mean distance of the 3 nearest neighbors of each cell was computed and averaged per section also allowing us to appraise the proximity of changes through time.

## Acknowledgments

We thank Antonella Rauseo and Juliette Cicchini-Bauquis for their help in animal handling, genotyping and immunofluorescence experiments, Fabrizio Thorel and Olivier Fazio from the transgenesis platform for their help in the generation of the transgenic knock-in line *Wnt5a:Tet-On3G* (Faculty of Medicine, University of Geneva), the animal facility team (Faculty of Medicine, University of Geneva), Jean-Pierre Aubry, Cécile Gameiro and Grégory Schneiter of the flow cytometry platform, the genomic platform and Nicolas Liaudet of the bioimaging platform (Faculty of Medicine, University of Geneva) for his substantial contribution with the counting and cell inter-distance analysis. We thank also the members of the Nef laboratory for helpful discussion and critical reading of the manuscript.

## Author Contributions

Conceptualization, H.A. and S.N.; Methodology: H.A. and B.C., Investigation, H.A.; Formal Analysis and Data Curation, H.A., I.S., C.M.R.; Writing – Original Draft, H.A. and S.N.; Funding Acquisition, S.N.; Resources, S.N.; Supervision, S.N.

## Supplementary figure legends

**Figure S1: Absence of a testicular phenotype in XX and XY** *Wnt5a:Tet-On3G* **embryos at E13.5**.

**Figure S2**: **Characterization of the urogenital tract of XX and XY** *Wnt5a:Tet-On3G* newborn pups.

**Figure S3**: **Absence of a reproductive phenotype in XX and XY** *Wnt5a:Tet-On3G* adult mice.

**Figure S4**: **Co-immunofluorescence for tdTOMATO and the progenitor markers NESTIN in E12.5 and E13.5 testis**.

**Figure S5: Comparison between** *Wnt5a* **and** *Tcf21* **expression in the developing testes**.

**Figure S6: Knock-in of the** *Tet-On 3G* **transgene into the mouse** *Wnt5a* **locus**.

**Figure S7: CRISPR/Cas9 sgRNA design and cloning**.

**Figure S8: Knock-in of the** *Tet-On 3G* **transgene into the murine** *Wnt5a* **locus**.

**Figure S9:** *Tet-ON 3G* **site of insertion in the genome in F1**.

