## Supplementary material for "Expression of *Wnt5a* defines the major progenitors of fetal and adult Leydig cells"

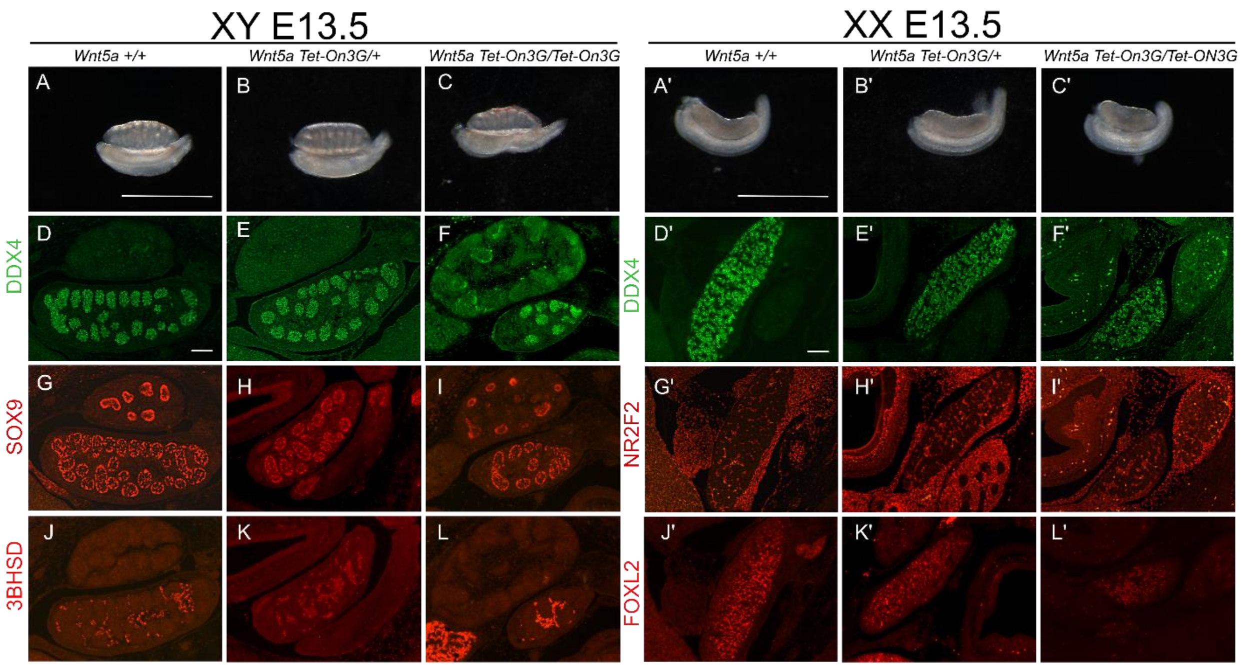


**Figure S1: Absence of a testicular phenotype in XX and XY *Wnt5a:Tet-On3G* embryos at E13.5.** The gonadal development of XY (A-L) and XX (A'-L') *Wnt5a* +/+ (control), *Wnt5a:Tet-On3G*/+ (heterozygous) and *Wnt5a:Tet-On3G*/*Wnt5a:Tet-On3G* (homozygous) embryos at E13.5 was evaluated by immunofluorescence for the germ cell marker (DDX4) (D-F and D'-F'), the Sertoli cell marker (SOX9) (G-I), the Leydig cell marker (3BHSD) (J-K), the interstitial progenitor marker NR2F2 (G'-I') and the granulosa cell marker (FOXL2) (J'-L'). N=3 animals tested per genotype. While no gonadal phenotype was observed in heterozygous animals compare to control animals, we observed a slight reduction in the size of the gonads in homozygous gonads. Three animals were analyzed per genotype. Scale bars: 1mm (A-C, A'-C'), 50 µm (D-L, D'-L').

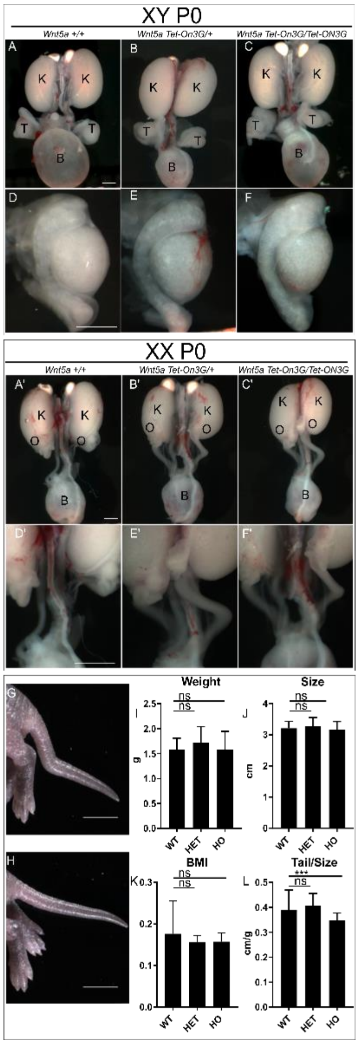


**Figure S2: Characterization of the urogenital tract of XX and XY Wnt5a:Tet-On3G newborn pups.** Evaluation of XY (A-F) and XX (A'-F') urogenital tract phenotype of *Wnt5a* +/+ (control), *Wnt5a:Tet-On3G*/+ (heterozygous) and *Wnt5a:Tet-On3G*/*Wnt5a:Tet-On3G* (homozygous) newborn pups (PO) (n=20). Photomicrographs of the male and female urogenital tract (A-C and A'-C') and gonads (D-F, D'-F') did not reveal any particular defects. (G-H) The tail of the homozygous *Wnt5a:Tet-On3G*/*Wnt5a:Tet-On3G* mice displays frequently a small bending. Evaluation of body weight (I), body length (J), body mass index (K) and tail/body length (L). ns, not significant; *** P <0.001. Abbreviations: T, testis; K, kidney; O, ovary; B, bladder. Scale bars: 1mm (A-F, A'-F'), 1mm (G-H).

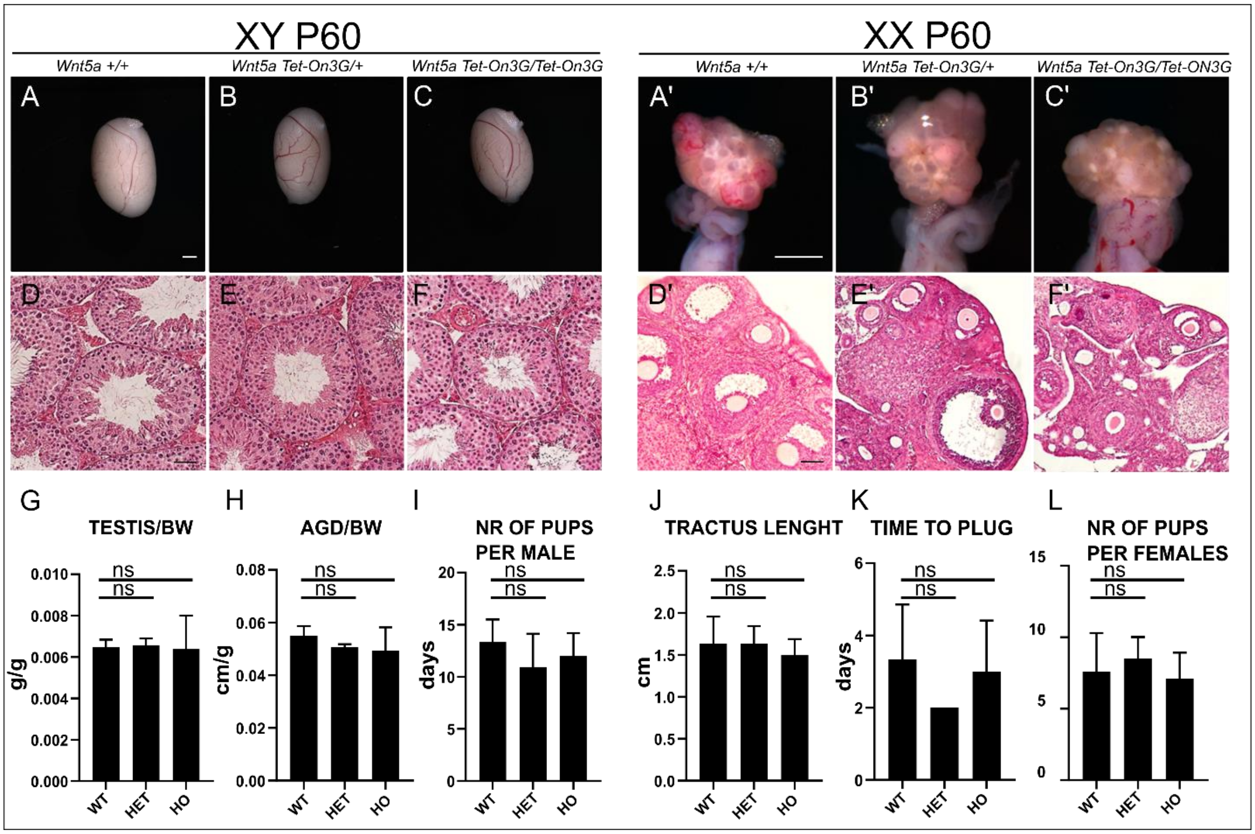


**Figure S3**: **Absence of a reproductive phenotype in XX and XY *Wnt5a:Tet-On3G* adult mice.** The testicular, ovarian and reproductive phenotype of *Wnt5a* +/+ (control), *Wnt5a:Tet-On3G*/+ (heterozygous) and *Wnt5a:Tet-On3G*/*Wnt5a:Tet-On3G* (homozygous) adult mice was evaluated at P120. No abnormalities was observed in the morphology (A-C, A'-C') and histology (D-F, D'-F') of adult testes (A-F) and ovaries (A'-F'). In males, testis weight (G), ano-genital distance (H) and number of animals per litter (I) are not affected by the genotype. In females, the length of the uterus (J), the time to plug (K) and the number of pups per female (L) is not affected by the genotype. Ns = not significant. N=4 per genotype. ns, not significant. Scale bars: 1mm (A-C, A'-C'), 10 μm (D-F, D'-F').

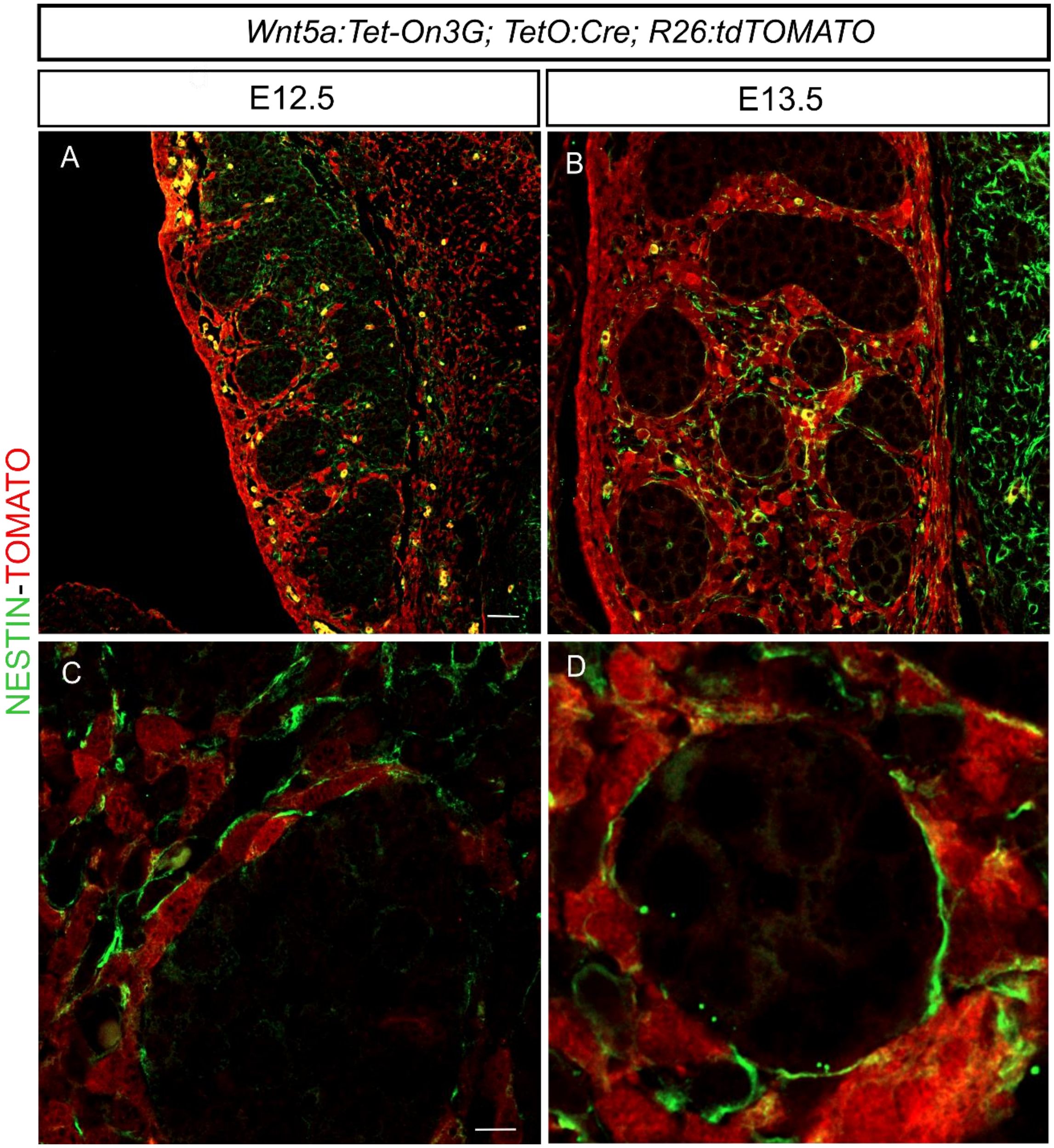


**Figure S4: Co-immunofluorescence for tdTOMATO and the progenitor markers NESTIN in E12.5 and E13.5 testis.** Evaluation of Tomato-labelled cells and NESTIN expressing cells in *Wnt5a:Tet-On3G;TetO:Cre;R26:tdTOMATO* embryos induced for 24 hours at E11.5. Testis were analyzed at E12.5 (A,C), and E13.5 (B,D) by co-immunofluorescence for tdTOMATO and the progenitor markers NESTIN. Scale bars: 20 µm (A,B), 5µm (C,D).

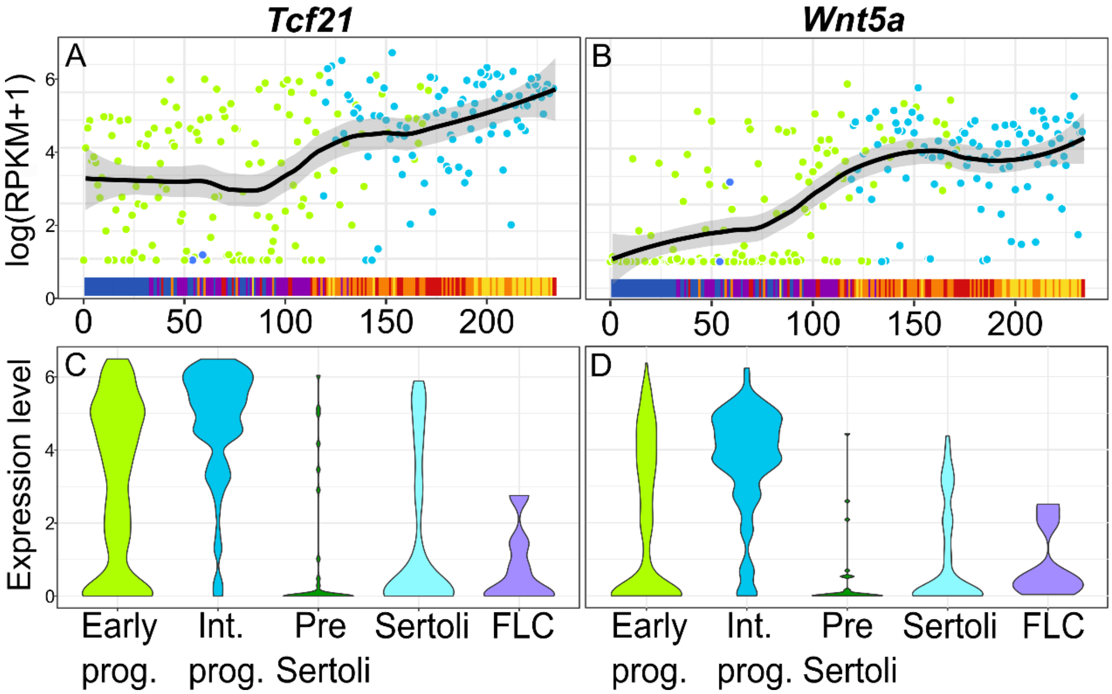


**Figure S5: Comparison between *Wnt5a* and *Tcf21* expression in the developing testes.** (A, B) Expression profiles of *Tcf21* (A) and *Wnt5a* (B) in NR5A1$^{+}$ somatic cell population of the fetal testis as determined by scRNA sequencing (Stévant et al 2018). The solid line represents the loess regression, and the fade band is the 95% confidence interval of the model. Dots represent single cells; the bar at the bottom of each graph represents the embryonic stages of single cells. (C, D) Violin graphs representing the single-cell expression profiles of *Tcf21* and *Wnt5a* in the different NR5A1$^{+}$ somatic cell population of the fetal testis. This profiling analysis includes 401 individual *Nr5a1*$^{+}$ transcriptomes from early progenitors (Early prog.), interstitial progenitors (Int. prog.), pre-Sertoli, Sertoli Cells and Fetal Leydig Cells (FLC). The width of the violin indicates frequency at that expression level. Note that *Tcf21* is expressed as early as E10.5 in multipotent early progenitors, while Wnt5a is restricted to the interstitial steroidogenic progenitors and starts to be expressed around E12.5.

**Supplementary material and methods**

**Generation of the *Wnt5a:TetOn3G* knock-in line**

We aim to introduce the Tetracycline inducible expression system transgene (*Tet-On 3G*) in the murine chromosome 14 *Wnt5a* locus, under the control of the *Wnt5a* endogenous promoter. CRISPR-Cas9 technology was used to genetically manipulate the mouse genome and introduce two double strand breaks in the sequence of the *Wnt5a* 5'UTR. For this purpose, two sgRNAs, here referred to as sgWnt5a ATG-1 and sgWnt5a1 3', were designed to cut 8 nucleotides (nt) after the ATG and 0.8 kb downstream the site of initiation of transcription. A targeting vector carrying the *Tet-On 3G* sequence was employed to promote transgene insertion into the genome through homology-directed repair (HDR) (**Figure S6**).

A

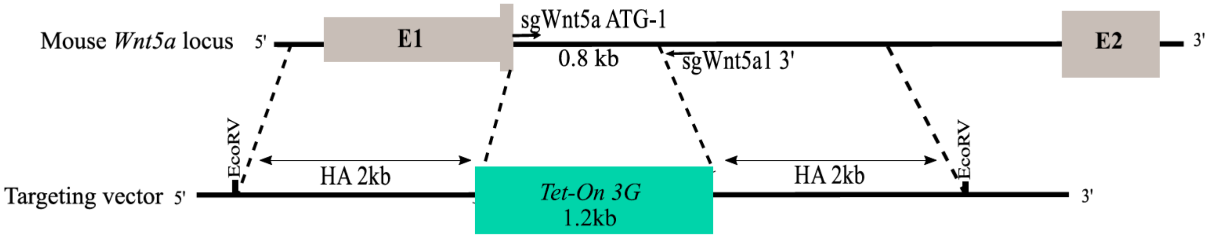


**Figure S6: Knock-in of the *Tet-On 3G* transgene into the mouse *Wnt5a* locus.** Diagram of knock-in generation strategy. A. sgWnt5a ATG-1, sgWnt5a1 3' site of cuts and target vector design are shown. The position of the 2kb homology arms (HA) in the target vector used to support HDR is indicated.

### Single guide RNAs design and cloning

The sgRNA targeting sequences were designed with a length of 20nt, corresponding to sequences at the extremities of the targeted 0.8kb fragment, followed by a 3nt PAM sequence (NGG). sgWnt5a ATG-1 and sgWnt5a1 3' were cloned in pX330-U6-Chimeric_BB-CBh-hSpCas9 (Addgene, Watertown, Massachusetts, USA). A guanine at the 5' end was inserted as $N_{21}$ since this was beneficial for the expression of the plasmid by the human U6 promoter (**Figure S7**). Putative off-target activities were evaluated with *'The CRISPR Software Matchmaker'*.

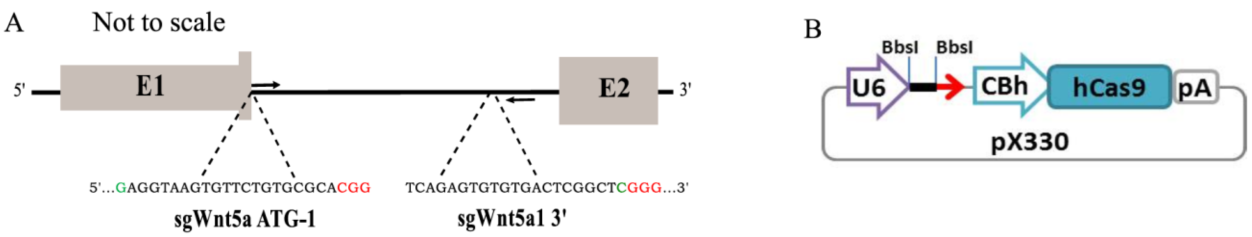


**Figure S7: CRISPR/Cas9 sgRNA design and cloning.** (A) Schematic diagram of the mouse targeted *Wnt5a* locus. The sgWnt5a ATG-1 and sgWnt5a1 3' sequences are shown upstream of the protospacer adjacent motif (PAM, in red); $N_{21}$ guanine and cytosine in green. (B) Map of the cloning plasmid used for zygote injection together with the restriction sites used for the insertion of the sgRNA sequences.

The sgRNAs (**Table S1**) were prepared as follows. Five µg of the vector were digested with BbsI (New England Biolabs, Ipswich, USA), gel purified with the QIAquick gel extraction kit (Qiagen, Hilden, Germany), and ligated with 600 ng of annealed forward and reverse sgRNAs using T4 ligase enzyme (New England Biolabs, Ipswich, USA). The ligated PX330-sgRNAs were transformed into Max Efficiency DH5α Competent Cells according to the manufacturer protocol (Invitrogen, Carlsbad, USA). Selection of pX330-sgRNA clones containing sgRNA sequences was performed by colony PCR with the primers PX330-screen (5' - GGCCTATTTCCCATGATTCC-3') and the antisense oligo of sgWnt5a ATG-1 and sgWnt5a1 3'.

| | |
|---|---|
| Wnt5a1 ATG 1 (S) | CACCGAGGTAAGTGTTCTGTGCGCA |
| Wnt5a1 ATG 1 (AS) | AAACTGCGCACAGAACACTTACCTC |
| Wnt5a1 3' 5 (S) | CACCGAGCCGAGTCACACACTCTGA |
| Wnt5a1 3' 5 (AS) | AAACTCAGAGTGTGTGACTCGGCTC |

**Table S1: sgRNA sequence list**. In red are the nucleotides complementary to BbsI sticky ends, $N_{21}$ guanine and cytosine in green.

**Targeting vector design**

The targeting vector sequence contained the Tetracycline inducible expression system transgene (*Tet-On 3G*), provided with a stop and poly(A)-addition signal. In order to promote homology directed repair, *Tet-On 3G* was flanked by 2kb arms, homologous to the *Wnt5a* wild type locus. The entire cassette was than inserted in pBluescrip-II-Ks between EcoRV sites. (Genetech, South San Francisco, California) (see **Figure S6**).

All plasmids were diluted in EmbryoMax® M2 Medium (Merk, Readington, USA) for pronuclear microinjection at concentration of respectively 2.5 ng/µl of sgWnt5a ATG-1, 2.5 ng/µl sgWnt5a1 3' and 5ng/µl recombination template. The three plasmids were injected in overall 479 mouse zygotes in two separate rounds of injections. Only 253 reached the blastocyst stage and were transferred into 6 foster mothers. Three living pups were obtained after the first injection and eleven pups after the second. Ultimately, 1/479 zygotes contained the desired mutation knocked in the Wnta 5’-UTR and was able to transmit it to the F1 generation.

**Identification and genotyping of *Wnt5a:TetOn3G* animal founders**

Injected embryos were genotyped using genomic DNA extracted from tail biopsies with DirectPCR Lysis Reagent (Viagen Biotech, Los Angeles, USA) and proteinase K. For PCR products larger than 1.5 kb, genomic DNA from the liver was extracted with DNeasy Blood & Tissue Kits (Qiagen) and Platinum SuperFi DNA Polymerase (Invitrogen, Carlsbad, California, USA) was used for the amplification, according to manufacturer protocol. The PCR primers used for genotyping are represented in **Figure S8** and their sequences listed in **Table S2**.

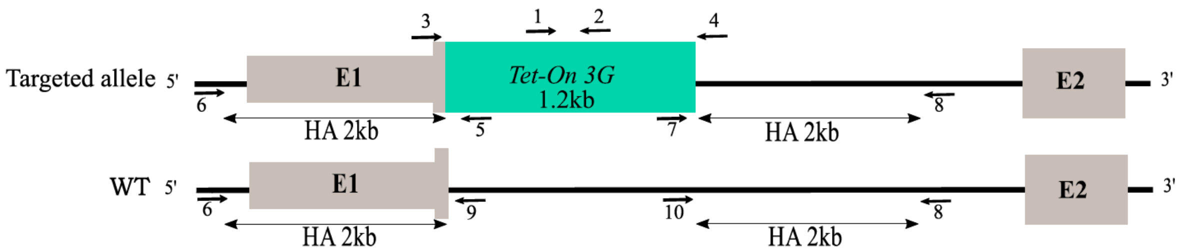


**Figure S8: Knock-in of the *Tet-On 3G* transgene into the murine *Wnt5a* locus.** Schematic map of the Wnt5a locus together with the locations of genotyping primers in the targeted and wild type alleles.

| | |
|---|---|
| GCTGCGTATTGGAGGAACAGG | forward 1 (21988) |
| TGCATTCTAGTTGTGGTTTGTCC | reverse 2 (21989) |
| CCGACGCTTCGCTTGAATTCC | forward 3 (21990) |
| GGAAATTGTTATCGAGGCGTGG | reverse 4 (21991) |
| AGTTTATGACTTTGCTCTTGTCC | reverse 5 (21997) |
| CTAGCAAACCTCTGACACCAGG | forward 6 (21992) |
| GGCCTCTTCATCGGGAATGC | forward 7 (21996) |
| AGAAGCCGGAGCATCTGTCG | reverse 8 (21995) |
| AAAAGTAGGGAGCGTCCGTGC | reverse 9 (22152) |
| TGGAGACTCTAGGGCCTCTG | forward 10 (22153) |

**Table S2: List of the primers used for genotyping.**

*Tet-ON 3G* cassette presence was confirmed in F0 animals with the primers 1-2 that anneal specifically in the *Tet-ON 3G* transgene and give an amplicon of 358 bp. Moreover, a second PCR with the couple of oligonucleotides 3-4 allowed us to distinguish putative homo and heterozygous animals. Subsequently the germline transmission of *Tet-ON 3G* insertion in F1 generation was confirmed by genotyping the offspring with the set of primers 1-2 and 3-4. F1 generation from founders F07, F013 and F014, which did transmit the *Tet-ON 3G,* were further investigated with primers from 5 to 10 (**Table S2**) that indicated the precise site of insertion in the genome: 9-6 **Figure S9A**, 2029 bp) and 5-6 **(Figure S9C** , 2018 bp), amplified the 5'-homology arm of respectively the wildtype (wt) and targeted allele. Correspondingly, 10-8 (**Figure S9B** 2349 bp) and 7-8 (**Figure S9D** 2281 bp) amplified the 3'-homology.

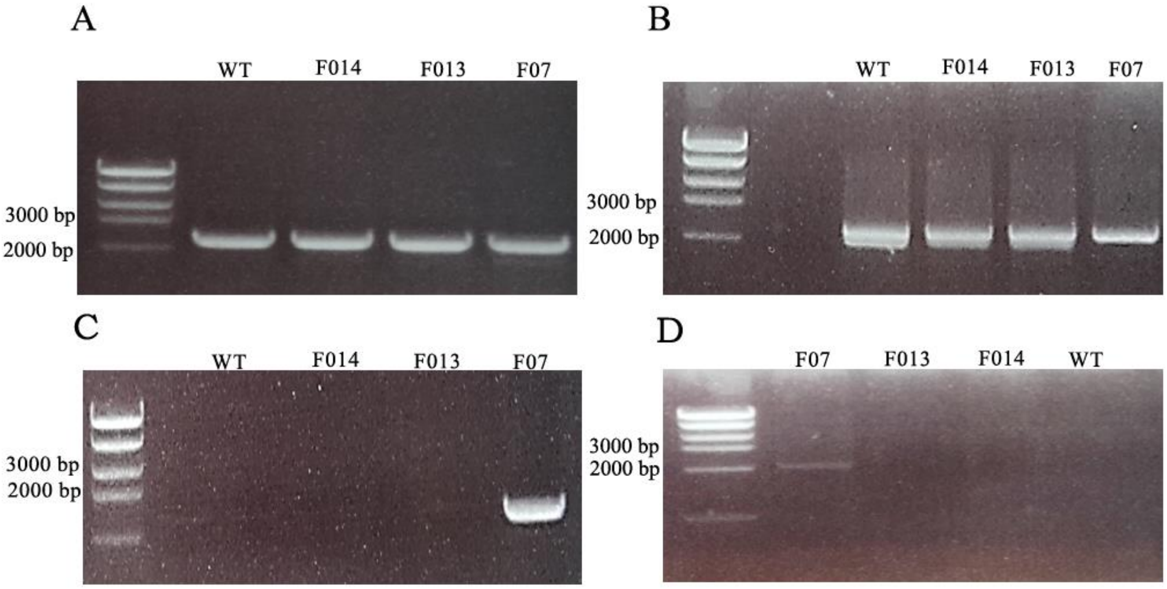


**Figure S9: *Tet-ON 3G* site of insertion in the genome in F1**. PCR amplifications spanning from intron 1 until the *Wnt5a* endogenous locus outside the homology arms show presence of a wt allele in all F1 animals (A-B). However, the amplicon *Tet-ON 3G*-chromosome 14 is present only in the F07 litters, giving evidence for a knock-in line, whereas for F013 and F014 we do not observe any amplification (C-D).

Animals positive for 1-2, 3-4, 5-6 and 7-8 were considered as carriers of a knock-in allele, those positive for 1-2, 3-4, 6-5, 7-8 as carriers of a transgenic insertion and finally mice positive only for 9-6 and 10-8 as wild type.

| | |
|---|---|
| CCGTAGCTCCAGCTTCACC | *TetO:Cre* 22075 |
| ACTTGCAGTTCTTGCAGGC | *TetO:Cre* 22076 |
| CATTTCGTGATGAATGCCAC | *TetO:Cre* 22077 |
| AAGGGAGCTGCAGTGGAGTA | *R26* wild type forward |
| CCGAAAATCTGTGGGAAGTC | *R26* wild type reverse |
| GGCATTAAAGCAGCGTATCC | *Tomato* mutant reverse |
| CTGTTCCTGTACGGCATGG | *Tomato* mutant forward |

**Table S3: List of primer pairs used for *TetO:Cre* and *R26:Tomato* loci genotyping.**

| | |
|---|---|
| TGAAGCAGGCCGTAGGACAG | *Wnt5a* forward |
| ACACTTACAGGCTACATCTGCCAG | *Wnt5a* reverse |
| CCTCAAAGTGGGCATGAGAC | *Nr2f2* forward |
| TGGGTAGGCTGGGTAGGAG | *Nr2f2* reverse |
| CTGTTTCACGCGGATGACT | *Amh* forward |
| CTCAGGCTCCAGGGACAG | *Amh* reverse |
| AAGTATGGCCCCATTTACAGG | *Cyp11A1* forward |
| TGGGGTCCACGATGTAAACT | *Cyp11A1* reverse |
| TTCCAGAAGACGCACTACCC | *Arx* forward |
| TCTGTCAGGTCCAGCCTCAT | *Arx* reverse |

**Table S4: List of primer pairs used for Real time PCR.**
